# Epithelial Glutamate Retention Protects against Colitis

**DOI:** 10.64898/2026.04.18.718419

**Authors:** Hideya Iwaki, Yunosuke Yasuda, Nobufumi Kato, Hiroshi Kitamura, Hidehiro Hayashi, Shohei Murakami, Hideyo Sato, Fan-Yan Wei, Shinji Fukuda, Tomoyoshi Soga, Takashi Kamei, Yoichi Kakuta, Atsushi Masamune, Hiroki Sekine, Hozumi Motohashi

**Author notes:** **Corresponding authors:** Hiroki Sekine, Ph.D., Department of Biochemistry and Molecular Biology, Graduate School of Medical Science, Yamagata University, Yamagata 990-9585, Japan, Hozumi Motohashi, M.D., Ph.D., Department of Medical Biochemistry, Graduate School of Medicine, Tohoku University, Sendai 980-8575, Japan.

## Abstract

Inflammatory bowel disease (IBD) is a chronic inflammatory disorder of the gastrointestinal tract that encompasses ulcerative colitis and Crohn’s disease. Here we identify the cystine/glutamate antiporter xCT as being markedly upregulated in the inflamed intestinal epithelium of patients with IBD. To clarify its functional contribution to disease pathogenesis, we performed genetic loss-of-function study and found that inhibition of xCT confers robust protection against dextran sulfate sodium (DSS)-induced colitis in mice. Intestinal epithelial cell (IEC)-specific deletion of xCT markedly attenuated colitis severity, demonstrating that epithelial xCT upregulation acts as a disease-exacerbating factor in IBD. Mechanistically, xCT deficiency preserved intracellular glutamate levels and protein polyglutamylation, thereby maintaining epithelial barrier integrity and protecting IECs from inflammatory injury. Consistently, pharmacological inhibition of glutamine synthetase, which increases intracellular glutamate, exerted a potent anti-inflammatory effect on the DSS-induced colitis. These findings identify intracellular glutamate retention in IECs as a previously unrecognized mechanism of epithelial protection and highlight both inhibition of xCT-dependent glutamate efflux and suppression of glutamine synthetase as potential therapeutic strategies for IBD.

## Introduction

Inflammatory bowel disease (IBD) is a chronic relapsing inflammatory disorder of the gastrointestinal tract and includes ulcerative colitis and Crohn’s disease. The global prevalence of IBD continues to rise, with disease onset most commonly occurring in early adulthood, necessitating lifelong medical support from a young age (Alatab et al., 2020; Cosnes et al., 2011). Current therapeutic strategies are largely based on immunosuppression, which broadly interferes with physiological immune functions and is associated with substantial long-term adverse effects (Matsuoka et al., 2018; Lamb et al., 2019). These limitations underscore an urgent need to alternative therapeutic approaches that ameliorate intestinal inflammation without compromising systemic immune competence.

The cystine/glutamate antiporter system xc□ comprises two subunits: the light chain xCT, encoded by *Slc7a11*, and the heavy chain 4F2hc, encoded by *Slc3a2* (Parker et al., 2021). xCT functions as the catalytic subunit of this heterodimer, mediating the uptake of extracellular cystine in exchange for intracellular glutamate. This transport activity directly links amino acid trafficking to cellular redox homeostasis, as imported cystine serves as a rate-limiting substrate for glutathione biosynthesis. Notably, sulfasalazine (SSZ), a frontline therapy for IBD, inhibits xCT activity and is widely used as a pharmacological tool to suppress system xc□ function in experimental settings (Gout et al., 2001). The clinical use of SSZ in IBD therefore raises the possibility that xCT-mediated transport contributes to disease pathophysiology; however, its precise functional role in the intestinal inflammatory context, and the downstream consequences of its inhibition, remain to be defined.

Using publicly available data sets, we found that SLC7A11 expression is markedly upregulated in the inflamed intestinal epithelium of patients with IBD, suggesting that elevated xCT activity has a modulatory role in the disease pathogenesis. To elucidate the contribution of xCT to IBD pathogenesis, we performed loss-of-function study using a dextran sulfate sodium (DSS)-induced colitis model in mice. We found that both genetic and pharmacological inhibition of xCT markedly attenuated colitis severity. Further mechanistic analyses revealed that intracellular glutamate retention in intestinal epithelial cells (IECs) promotes resistance to inflammatory injury by preserving epithelial barrier integrity. These findings implicate epithelial amino acid homeostasis as a previously underappreciated determinant of intestinal inflammation and establish intracellular glutamate retention in IECs as a conceptual framework for understanding epithelial resilience in IBD.

## Results

### xCT knockout mice are resistant to DSS colitis

By re-analyses of publicly available data sets (Czarnewski et al., 2019; Argmann et al., 2023), we found that *SLC7A11* expression is markedly upregulated in the inflamed intestinal epithelium of patients with IBD, both ulcerative colitis and Crohn’s disease (Figure 1A). Similar increase in the expression of *Slc7a11* was also observed in the DSS-induced colitic in mice (Figure 1B), which was verified in our experimental setting (Figure 1C).

**Fig. 1.**
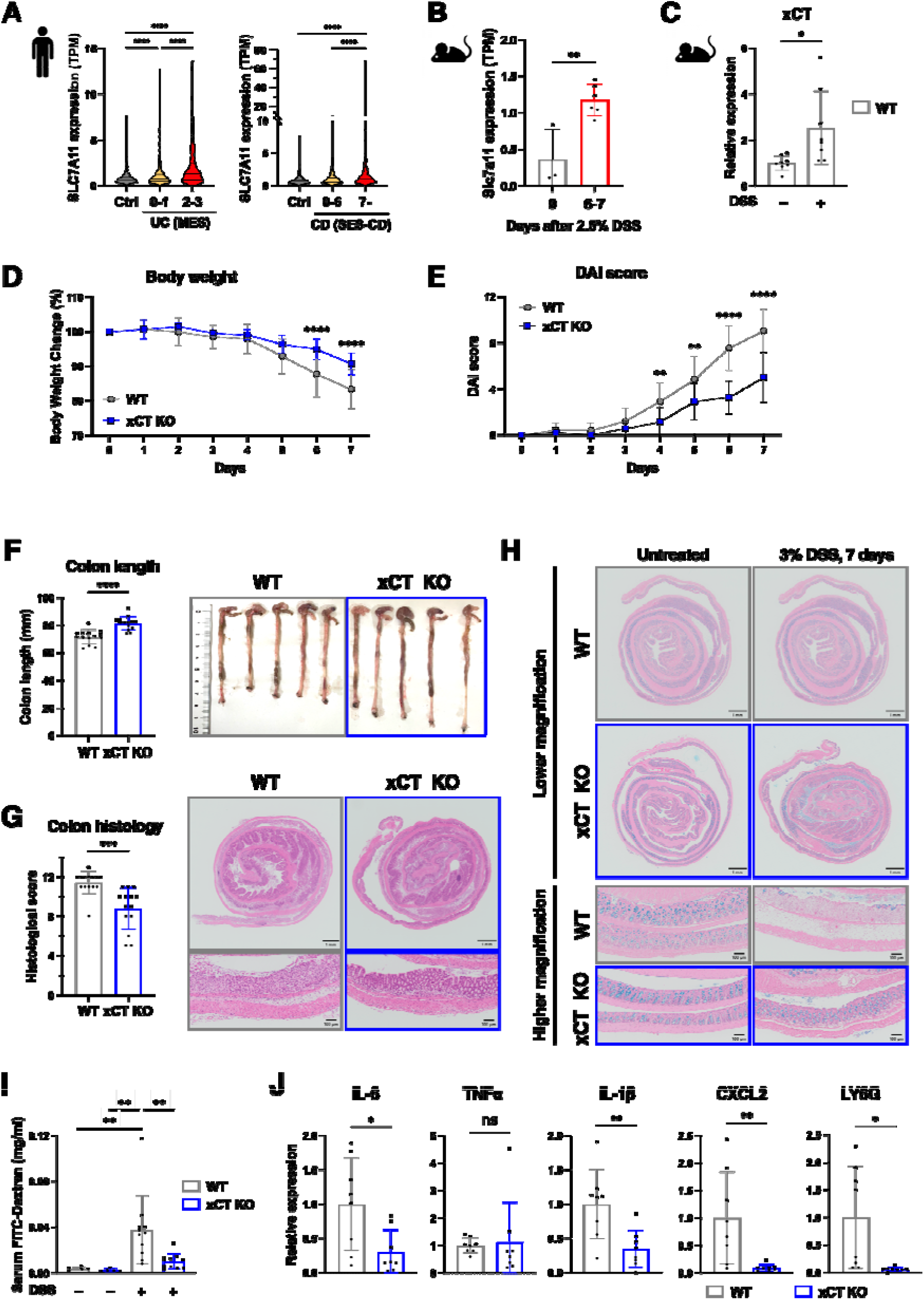
xCT deficiency shows anti-inflammatory effects on DSS colitis. **A.** mRNA Expression of *SLC7A11*, encoding xCT, in human inflammatory bowel diseases. Publicly available data sets (GSE186507, GSE193677, GSE206285, GSE207465) were used for analysis. UC: ulcerative colitis, MES: Mayo endoscopic subscore, CD: Chrohn’s disease, SES-CD: Simple endoscopic score for CD. **B.** mRNA Expression of *Slc7a11*, encoding xCT, in mouse colitic model induced by 2.5% DSS administration. Publicly available data sets (GSE131032) were used for analysis. **C.** RT-PCR for measuring mRNA expression of xCT in colon tissues of wild-type (WT) mice with or without 3% DSS treatment on day 6 (DSS – WT mice; n=8, DSS + WT mice; n=8). β-Actin was employed for normalization. Mean values of DSS – WT mice were set as 1. **D-G.** WT and xCT KO mice were treated with 3% DSS for 7 days and sacrificed on day 7 for analysis (n=14 per group). Body weight changes **(D)**, DAI score **(E)**, and colon length measurement and macroscopic observation of colon tissues **(F)** are shown. Histological score and HE staining of colon tissues are shown with scale bars corresponding to 1 mm and 100 μm for lower and higher magnifications, respectively (**G**). Data are shown as mean□±□SD from two independent experiments. **H.** Alcian blue staining of colon tissues from the mice sacrificed on day 7 after 3% DDS administration for detecting mucus. Untreated samples were obtained from mice given tap water. Representative micrographs are shown (n=2 per group). Scale bars correspond to 1 mm for lower magnification and 100 μm for higher magnification. **I.** Evaluation of intestinal permeability by FITC-dextran. FITC-dextran was quantified in serum with or without 3% DSS treatment on day 3 (DSS – WT mice; n=5, DSS – xCT KO mice; n=5, DSS + WT mice; n=10, DSS + xCT KO mice; n=11). Data are shown as mean□±□SD from three independent experiments. **J.** RT-PCR for measuring mRNA expression of cytokines, chemokines and a neutrophil marker, Ly6G, in colon tissues with 3% DSS treatment on day 6 (WT mice; n=8, xCT KO mice; n=8). β-Actin was employed for normalization. Mean values of WT mice were set as 1. Data are shown as mean□±□SD from two independent experiments. Data represent means ± standard deviation. **A**, **B**, and **I** were analyzed by one-way ANOVA. **D** and **E** were analyzed by two-way ANOVA. **C**, **F**, **G**, and **J** were analyzed by two-sided Student’s t-test. *p<0.05, **p<0.01, ****p<0.0001; ns, not significant.

Expecting that elevated xCT activity has a modulatory role in the disease pathogenesis, we first assessed *Slc7a11*^−/−^ (xCT KO) mice in DSS colitis (Sato et al., 2005). Although the precise mechanism of its colitogenicity remains incompletely understood, DSS, a sulfated polysaccharide, is thought to be toxic to intestinal epithelial cells (IECs), compromising barrier integrity and leading to inflammatory immune response to various luminal antigens (Kiesler et al., 2015; Wirtz et al., 2017). xCT KO mice were found more resistant than wild-type (WT) control mice, exhibiting significantly less body weight loss and disease activity index (DAI) score (Figure 1D and 1E). DSS colitis causes shortening of the intestinal tract, which reflects the severity of colitis (Chassaing et al., 2014). The shortening of colon length was attenuated in xCT KO mice compared with WT mice (Figure 1F). Histological observation on day 7 after the initiation of DSS treatment showed that the colon epithelia of WT mice were severely damaged with heavy infiltration of inflammatory cells, whereas intestinal glands were well preserved in the colon epithelia of xCT KO mice (Figure 1G). Mucus secreting cells in the intestinal glands, which were detected by alcian blue staining, were dramatically decreased in WT mice but well retained in xCT KO mice (Figure 1H). These results suggest that whole animal xCT deficiency has an anti-inflammatory effect on DSS colitis.

To further characterize the alleviation of colitis in xCT KO mice, a barrier function was examined by the FITC-dextran administration experiment (Dawson et al., 2009). Whereas leakage of FITC-dextran to the serum was almost undetectable before the DSS treatment irrespective of the genotypes, the barrier function was significantly impaired in WT mice but well retained in xCT KO mice after 3 days of DSS administration (Figure 1I). Genes encoding proinflammatory cytokines IL-6 and IL-1β were suppressed in xCT KO colon compared with WT colon (Figure 1J). Expression levels of CXCL2, a neutrophil-attracting chemokine, and Ly6G, a neutrophil marker, were also reduced in xCT KO mice. These results suggest that the intestinal epithelial barrier function is well preserved and that neutrophil infiltration and other inflammatory events are attenuated in xCT KO mice.

### xCT deficiency in IECs is responsible for the resistance to DSS colitis

Although precise etiology of IBD is not well understood, immune system, gut microbiota and IECs are currently regarded as three major factors underlying IBD in human as well as DSS colitis in mice (Eichele et al., 2017) (Fig. 2A). We first conducted bone marrow transplantation (BMT) experiment to examine a contribution of xCT deficiency in immune cells to the pathogenesis of DSS colitis (Chang et al., 2020) (Figure S1A). Whole bone marrow cells of xCT KO mice and control WT mice in CD45.1 background were transplanted to lethally irradiated WT recipient mice in CD45.2 background. The transplantation efficiencies were comparable between the two genotypes at 13-15 weeks after transplantation (Figure S1B). The DSS treatment induced colitis in both groups, recipient mice with WT bone marrow cells and those with xCT bone marrow cells. Their severity did not differ significantly different in terms of body weight loss, colon length shortening, histological changes, and inflammation-related gene expression (Figure S1C-S1F). Therefore, xCT deficiency in immune cells of bone marrow origins do not make any substantial contributions to the alleviation of DSS colitis.

**Fig. 2.**
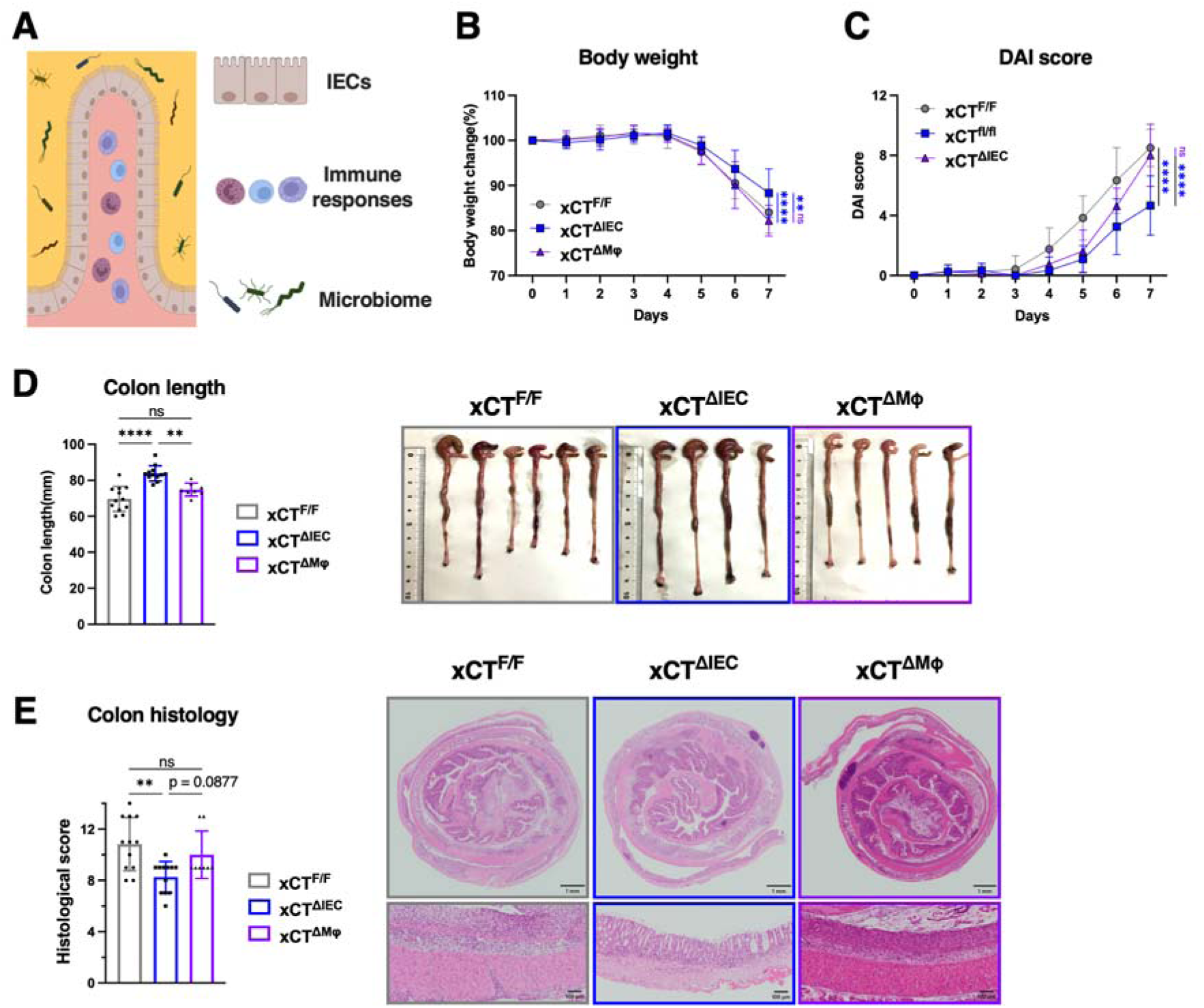
IEC-specific xCT disruption is sufficient to show anti-inflammatory effects on DSS colitis. **A.** Three major factors that play important roles in IBD and DSS colitis. **B-E.** *Slc7a11* floxed (xCT^F/F^) mice, *Slc7a11*^F/F^:Villin-Cre (xCT^ΔIEC^) mice and *Slc7a11*^F/F^:LysM-Cre (xCT^ΔMφ^) mice were treated with 3% DSS for 7 days and sacrificed for analysis on day 7 (xCT^F/F^ mice; n=14, xCT^ΔIEC^ mice; n=14, xCT^ΔMφ^ mice; n=8). Body weight changes (**B**), DAI score (**C**) and colon length measurement and macroscopic observation of colon tissues (**D**) are shown. Histological score and HE staining of colon tissues are shown with scale bars corresponding to 1 mm for lower magnification and 100 μm for higher magnification (**E**). Data are shown as mean□±□SD from four independent experiments. Data represent means ± standard deviation. **B** and **C** were analyzed by two-way ANOVA. **D** and **E** were analyzed by one-way ANOVA. **p<0.01, ****p<0.0001; ns, not significant.

We next examined whether whole animal xCT deficiency caused any alterations in gut microbiota, which might have elicited protective immune response. To ensure the same initial gut colonization, pregnant xCT KO mice were co-housed with pregnant WT mice, and their pups were bred in the same cage until 4 weeks after birth (Wirtz et al., 2017; Laukens et al., 2016; Brinkman et al., 2013) (Figure S2A). After weaning, the mice were separated according to the genotype and bred for additional 12-14 weeks to allow the gut microbiota to evolve and adapt to each host intestinal environment. Fecal samples before the colitis induction were subjected to 16S rRNA gene sequencing and metabolome analysis (Nagai et al., 2023; Forster et al., 2022). At the order level, a proportion of Bacillales, which belongs to the phylum Firmicutes, was higher in xCT KO mice than WT mice (Figure S2B). At the genus level, the proportions of Lachnospiraceae fcs020 group and Clostridium sensu stricto 1 were increased in xCT KO mice (Figure S2C). Firmicutes are known to synthesize short-chain fatty acids (SCFAs), which have been shown to possess anti-inflammatory effects in IBD and DSS colitis (Parada Venegas et al., 2019; Louis et al., 2017). However, the SCFAs, such as lactate, butyrate, and propionate, were not increased in the gut metabolites of xCT KO mice compared with those in WT mice (Figure S2D). Nevertheless, the xCT KO mice that were bred in the co-housing and co-breeding protocol (Figure S2A) were apparently resistant to the DSS colitis compared with WT mice (Figure S2E-S2G). These results suggest that modest microbiota changes observed in xCT KO mice are unlikely to explain their resistance to the colitis.

Finally, we investigated the contribution of IECs to the DSS colitis by exploiting xCT conditional knockout mice. We performed xCT deletion in IECs by crossing a mouse line harboring floxed alleles of *Slc7a11* (*Slc7a11*^F/F^; xCT^F/F^) with Villin-Cre transgenic mice line to generate IEC-specific *Slc7a11* knockout (*Slc7a11*^F/F^:Villin-Cre; xCT^ΔIEC^) mice (Madison et al., 2002). We also crossed *Slc7a11*^F/F^ mice with LysM-Cre mice to generate myeloid cell-specific *Slc7a11* knockout (*Slc7a11*^F/F^:LysM-Cre; xCT^ΔMφ^) mice to examine the xCT contribution in tissue-resident macrophages, which were not covered in the BMT experiment (Pan et al., 2022). DSS treatment was performed on xCT^ΔIEC^ mice, xCT^ΔMφ^ mice, and xCT^F/F^ mice as a control. In xCT^ΔIEC^ mice compared to xCT^F/F^ mice, body weight loss, DAI score, colon length shortening, and histological score were all significantly suppressed (Figure 2B-2E). In contrast, the colitis severity of xCT^ΔMφ^ mice was almost comparable to that of xCT^F/F^ mice. Therefore, we concluded that IEC is a major contributor to the anti-inflammatory effect of xCT deficiency conferring the resistance to the DSS colitis.

### xCT deficiency preserves protein polyglutamylation during DSS-induced colitis

To decipher a mechanism by which xCT inhibition in IECs protects colon tissues from inflammation, we first examined glutathione (GSH) levels in colon tissues before and after DSS treatment. Because xCT mediates cystine uptake for GSH synthesis, we anticipated that xCT deficiency might decrease GSH levels during DSS colitis. However, GSH levels were not altered in xCT KO mice compared with WT mice, either at baseline or following DSS treatment (Figure 3A) implying the activation of compensatory mechanisms for cysteine supply, such as de novo synthesis via the methionine transsulfuration pathway and/or uptake though alternative cysteine transporters, including ASC family transporters.

**Fig. 3.**
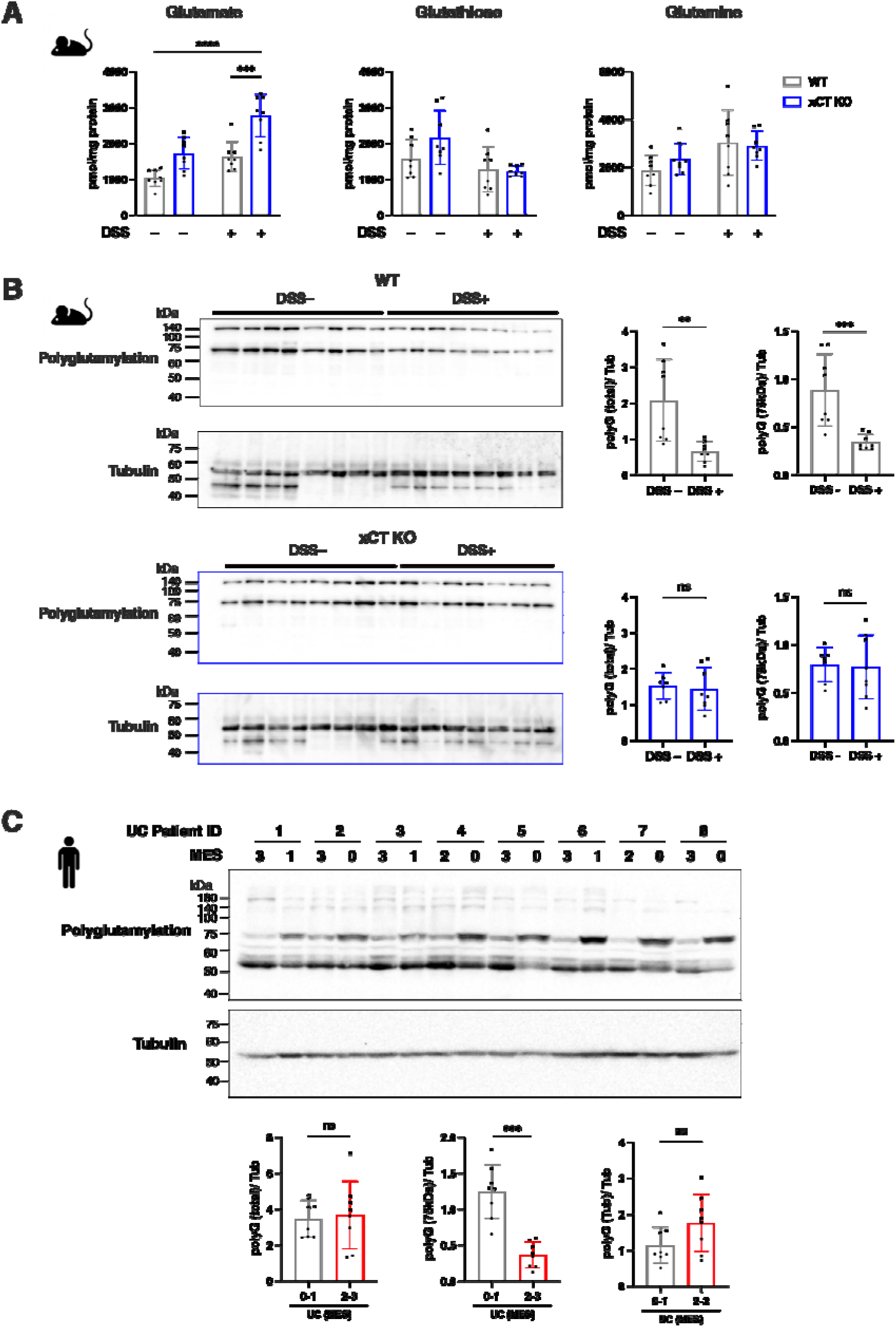
xCT deficiency retains intestinal glutamate and protein polyglutamylation during DSS colitis. **A.** Amino acid quantification in colon tissues of WT mice and xCT KO mice on days 0, 3, and 5 of 3% DSS treatment (n=3-4, per group). **B.** Protein polyglutamylation in colon epithelial tissues of WT mice and xCT KO mice with and without 3% DSS treatment on day 7. Tubulin was detected as a loading control. Band intensities were quantified using the Image J. **C.** Protein polyglutamylation in colon biopsy specimens of UC patients. The biopsy specimens during the active phase (MES3) and remission phase (MES1) were obtained. Tubulin was detected as a loading control. Band intensities were quantified using the Image J. Data represent means ± standard deviation. **A** was analyzed by one-way ANOVA. **B** and **C** were analyzed by non-parametric Mann-Whitney test. **p<0.01, ***p<0.001, ****p<0.0001; ns, not significant.

In contrast, glutamate levels were significantly elevated in the colon tissues of xCT KO mice compared with WT mice, but only following DSS treatment (Figure 3A). Considering a previous report describing reduced mucosal glutamate levels during DSS colitis (Shiomi et al., 2011), this observation suggestes that xCT uprgeulation during DSS colitis promotes glutamate export from IECs, whereas xCT deficiency prevents this loss and thereby facilitates intracellular glutamate retention.

Because glutamate serves as an essential substrate for protein polyglutamylation, we next assessed polyglutamylation levels in colon epithelial tissues of WT and xCT KO mice before and after DSS treatment (Figure 3B). DSS treatment reduced global protein polyglutamylation in WT mice, whereas this reduction was not observed in xCT KO mice. Consistently, analysis of colonic biopsy samples from patients with ulcerative colitis revealed a pronounced decrease in polyglutamylation of a ∼75 kDa protein in the disease-active state compared with the inactive state (Figure 3C). Notably, DSS-induced loss of polyglutamylation of this ∼75 kDa protein was evident in WT mice but was preserved in xCT KO mice (Figure 3B), closely mirroring the human disease pattern. Thus, these findings indicate that xCT deficiency promotes intracellular glutamate retention in IECs during colitis, thereby preserving protein polyglutamylation under inflammatory conditions.

### Reduced polyglutamylation in IECs compromises barrier function and enhances inflammatory responses

To investigate the functional consequences of glutamate retention and associated protein polyglutamylation in IECs, we employed Caco-2 cells as an *in vitro* IEC model. We first verified that culture in glutamine-free medium decreases global polyglutamylation in Caco-2 cells (Figure 4A), indicating that extracellular glutamine is a major source of intracellular glutamate under this experimental setting. We next assessed transepithelial electrical resistance (TEER), a quantitative measure of intestinal epithelial barrier integrity. Caco-2 cells cultured in glutamine-free medium, conditions under which polyglutamylation is attenuated, exhibited a significant reduction in TEER (Figure 4B), suggesting that epithelial barrier function is compromised under glutamine-restricted conditions. We also examined TNF-α–induced IL-8 expression, a hallmark inflammatory response in intestinal epithelial cells (IECs) relevant to IBD (Van De Walle et al., 2010; Friedrich et al., 2014). Glutamine deprivation enhanced TNF-α-induced IL-8 expression, and this effect was partially reversed by the xCT inhibitor SSZ, which promotes intracellular glutamate retention (Figure 4C). These findings support a role for intracellular glutamate in attenuating epithelial inflammatory responses.

**Fig. 4.**
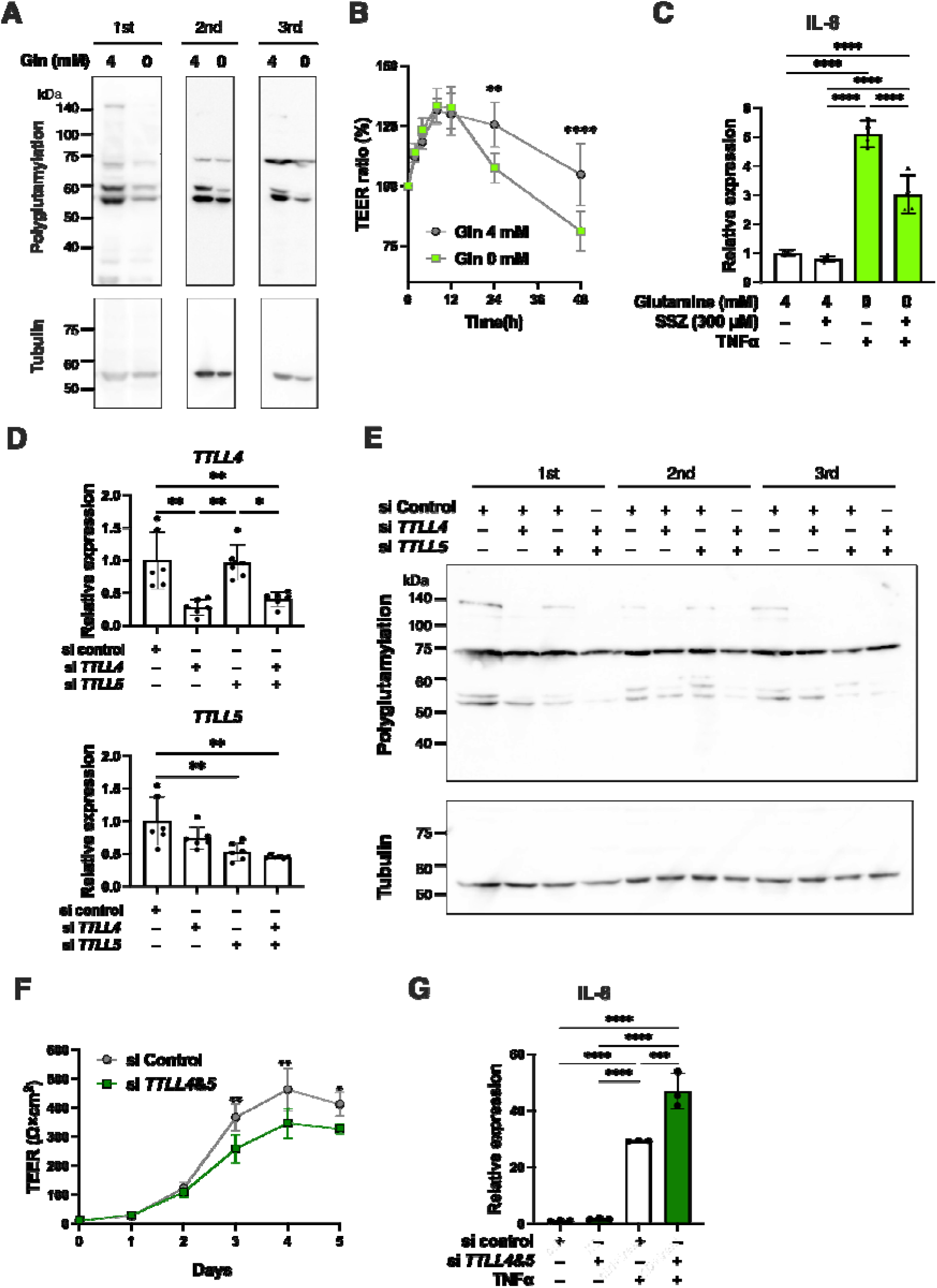
Inhibition of protein polyglutamylation compromises epithelial barrier function. **A.** Detection of polyglutamylated proteins with or without glutamine in Caco-2 cells. Tubulin was detected as a loading control. Results from three independent experiments are shown. **B.** Measurement of transepithelial electrical resistance (TEER) of Caco-2 cell culture with or without glutamine. Data are shown as mean□±□SD from four independent experiments. **C.** RT-PCR to measure mRNA expression of IL-8 in HCT116 cells following TNFα stimulation with or without glutamine and SSZ (300 μM) (n=4, per group). β-Actin was employed for normalization. A mean value of samples with 4 mM glutamine without SSZ was set as one. Data are shown as mean□±□SD from four independent experiments. **D.** Knockdown efficiency of *TTLL4* and *TTLL5* in Caco-2 cells. Data are shown as mean□±□SD from six independent experiments. **E.** Detection of polyglutamylated proteins with or without knockdown of *TTLL4* and/or *TTLL5* in Caco-2 cells. Tubulin was detected as a loading control. Results from three independent experiments are shown. **F.** Measurement of TEER of Caco-2 cell culture with or without *TTLL4* and *TTLL5* double knockdown. Data are shown as mean ± SD from four independent experiments performed in duplicate. **G.** RT-PCR to measure mRNA expression of IL-8 in Caco-2 cells following TNFα stimulation with or without knockdown of *TTLL4* and *TTLL5* (n=3, per group). β-Actin was employed for normalization. A mean value of control siRNA-treated samples without TNFα stimulation was set as one. A representative result is shown from two independent experiments. Data represent means ± standard deviation. **B** and **F** were analyzed by two-way ANOVA. **C**, **D** and **G** were analyzed by one-way ANOVA. * p<0.05, ** p<0.01, *** p<0.001, **** p<0.0001.

To directly assess the contribution of protein polyglutamylation to epithelial barrier function and inflammatory response, we inhibited tubulin tyrosine ligase-like (TTLL) family enzymes, which catalyze protein polyglutamylation (Torrino et al., 2021). Combined knockdown of *TTLL4* and *TTLL5* in Caco-2 cells attenuated global polyglutamylation (Figure 4D and 4E), which resulted in decreased TEER (Figure 4F) and increased IL-8 expression in response to TNF-α (Figure 4G). Collectively, these findings indicate that protein polyglutamylation is a critical determinant of intestinal epithelial barrier integrity and inflammatory responsiveness. They further suggest that the protective effect of xCT inhibition in DSS-induced colitis arises, at least in part, from the prevention of inflammation-associated glutamate loss from IECs, thereby maintaining protein polyglutamylation, preserving epithelial barrier function, and limiting pro-inflammatory cytokine expression.

### Glutamine synthetase inhibition attenuates DSS-induced colitis

An emerging concept from our findings is that elevated intracellular glutamate levels in IECs exert anti-inflammatory effects during DSS-induced colitis. To further test this concept using an independent approach, we sought to increase intracellular glutamate by modulating glutamine-glutamate interconversion. Glutamine synthetase (GS) catalyzes the conversion of glutamate to glutamine in an ATP-dependent manner. Based on previous work demonstrating that administration of methionine sulfoximine (MSO), a pharmacological inhibitor of GS, increases glutamate levels in both serum and tissues (Villar et al., 2023), we evaluated the impact of GS inhibition on DSS colitis (Figure 5A).

**Fig. 5.**
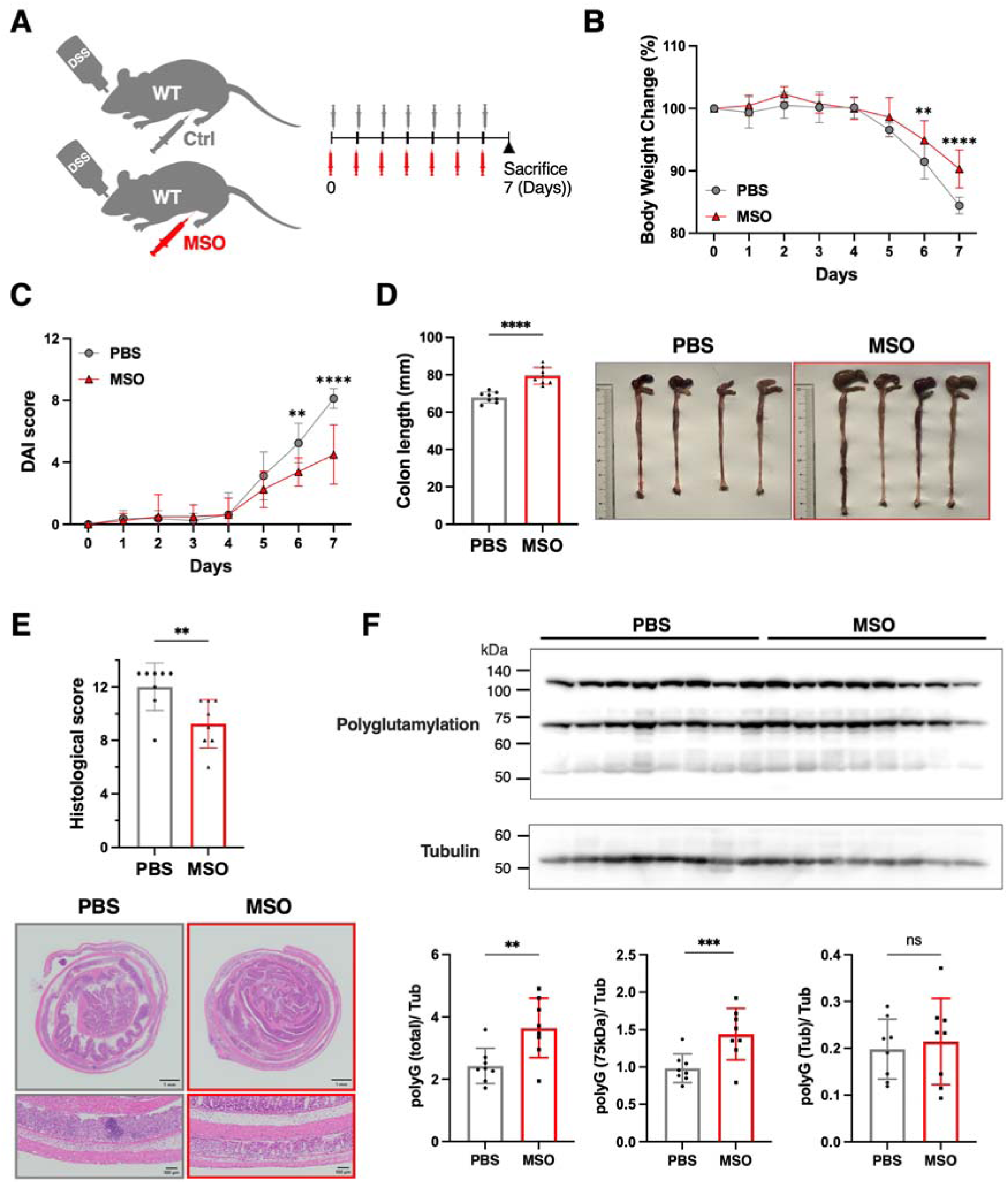
A glutamine synthetase inhibitor MSO shows anti-inflammatory effects on DSS colitis.A. Scheme of MSO administration experiment in 3% DSS colitis. WT mice were divided into two groups, MSO group (10 mg/kg/day, n=8) and control group (equal volume of PBS, n=8), and administered with MSO or PBS, respectively, by intraperitoneal injection once daily from day 0 to day 6. **B-E.** Effects of MSO on DSS colitis were examined. Body weight changes (**B**), DAI score (**C**), and colon length measurement and macroscopic observation of colon tissues (**D**) are shown. Histological score and HE staining of colon tissues are shown with scale bars corresponding to 1 mm for lower magnification and 100 μm for higher magnification (**E**). Data are shown as mean□±□SD from three independent experiments. **F.** Protein polyglutamylation in colon tissues of WT mice treated with MSO or PBS together with 3% DSS on day 5. Tubulin was detected as a loading control. Band intensities were quantified using the Image J. Data represent means ± standard deviation. **B** and **C** were analyzed by two-way ANOVA. **D-F** were analyzed by two-sided Student’s t-test. **p<0.01, ****p<0.0001; ns, not significant.

MSO administration significantly attenuated DSS-induced body weight loss, disease activity index (DAI), colon shortening, and histological damage in wild-type (WT) mice (Figure 5B-5E). These findings provide independent support for the notion that intracellular glutamate accumulation in IECs plays a protective role in colitis and contributes to resistance against DSS-induced intestinal inflammation.

### Pharmacological inhibition of xCT ameliorates DSS-induced colitis

Because prolonged systemic inhibition of GS may cause detrimental side effects, as suggested by the phenotypes observed in patients with congenital GS mutations (Häberle et al., 2005) and in GS knockout mice (He et al., 2010; Hakvoort et al., 2017), we reasoned that inhibition of xCT would represent a more suitable strategy to induce glutamate retention in intestinal epithelial cells when systemic administration is undertaken. We therefore employed pharmacological inhibition of xCT and examined whether this approach is sufficient to alleviate DSS-induced colitis in WT mice. Treatment with imidazole ketone erastin (IKE), a selective xCT inhibitor, resulted in marked attenuation of DSS-induced body weight loss, DAI score, colon length shortening, and histological injury (Figure 6A-6E).

**Fig. 6.**
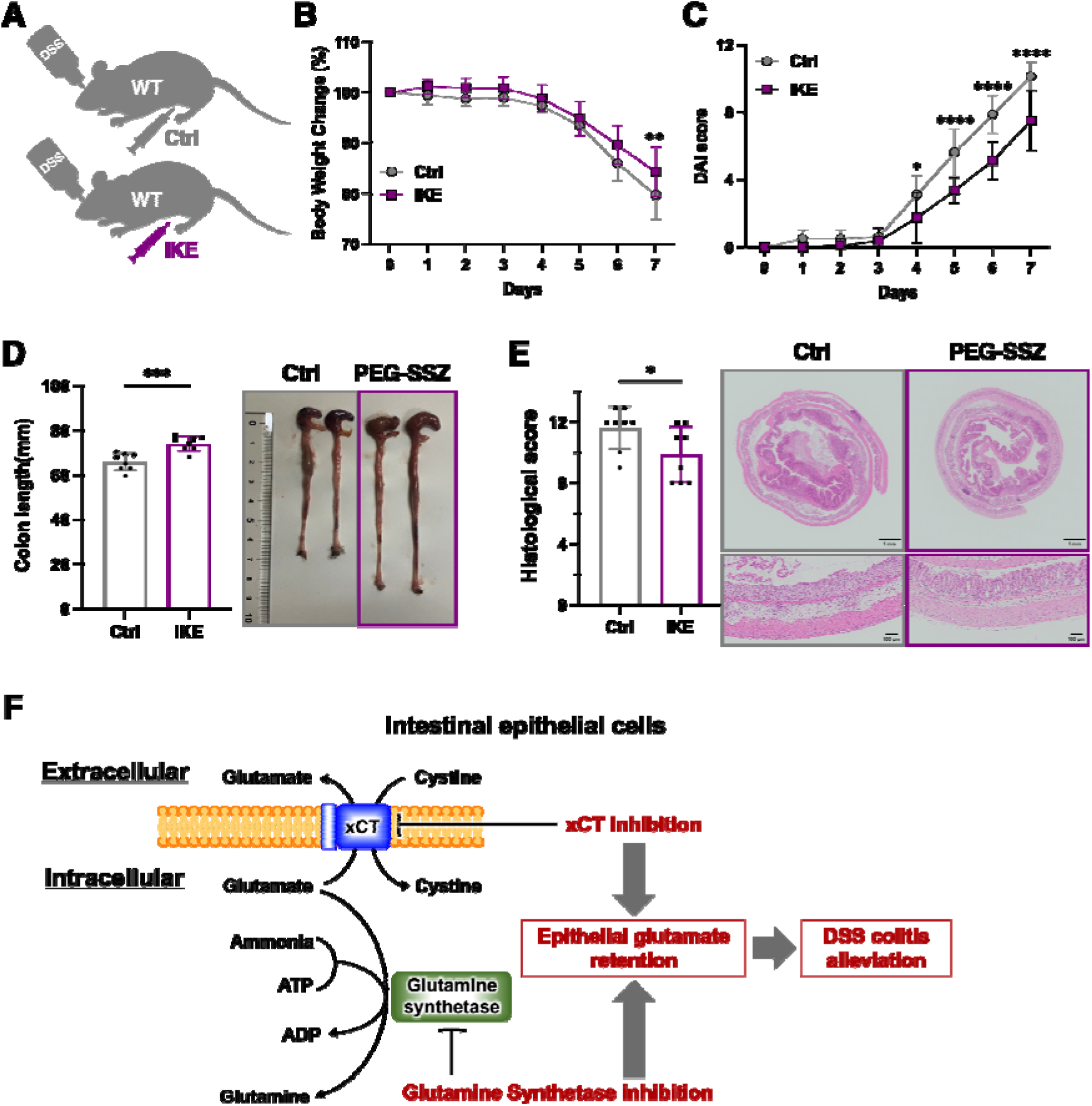
Inhibitors of xCT show anti-inflammatory effects on DSS colitis. **A.** Scheme of IKE administration experiment in 3% DSS colitis. WT mice were divided into two groups, IKE group (25 mg/kg/day, n=8) and control group (equal volume of the vehicle, n=8), and administered with IKE and the vehicle, respectively, by intraperitoneal injection once daily from day 0 to day 6. **B-E.** Effects of IKE on DSS colitis were examined. Body weight changes (**B**), DAI score (**C**), and colon length measurement and macroscopic observation of colon tissues (**D**) are shown. Histological score and HE staining of colon tissues are shown with scale bars corresponding to 1 mm for lower magnification and 50 μm for higher magnification (**E**). Data are shown as meanLJ±LJSD from three independent experiments. **F.** Two different approaches for promoting glutamate retention in intestinal epithelial cells. Both results in alleviation of DSS colitis. Data represent means ± standard deviation. **B** and **C** were analyzed by two-way ANOVA. **D** and **E** were analyzed by two-sided Student’s t-test. *p<0.05, **p<0.01, ***p<0.001, ****p<0.0001.

Together with the protective effect of MSO, these results demonstrate that pharmacological inhibition of either GS or xCT effectively suppresses DSS-induced colitis (Figure 6F). They further suggest that modulation of epithelial amino acid metabolism may represent a therapeutic strategy that complements existing immunosuppressive treatments, potentially enabling dose reduction and mitigation of systemic side effects.

## Discussion

We found that xCT inhibition shows a potent anti-inflammatory effect on DSS colitis and that inhibition of xCT in IECs is responsible for protection from the colitis. Our data suggest that resistance to the DSS colitis stems from the glutamate retention in IECs, which enhances protein polyglutamylation allowing increase of intestinal epithelial barrier function and decrease of cytokine production by IECs. In support of this notion, administration of a GS inhibitor, which is supposed to maintain intracellular glutamate levels, has an anti-inflammatory effect on DSS colitis. An important concept that emerged from this study is that IECs can be a therapeutic target and that glutamate is critical for the robustness of IECs.

Protein polyglutamylation is a reversible post-translational modification in which one or more glutamate residues are added as lateral chains to glutamate residues within target proteins (Ruse et al., 2022). Although this modification has been extensively characterized on α- and β-tubulin, where it regulates microtubule dynamics and associated protein interactions (Eddé et al., 1990; Rüdiger et al., 1992), our analyses of human and mouse intestinal epithelial tissues revealed a prominent polyglutamylated species of approximately 75 kDa, substantially larger than tubulin. This finding suggests that a non-tubulin protein is a major target of polyglutamylation in IECs. Notably, in human colonic tissues, the polyglutamylation status of this 75 kDa protein showed a strong association with disease activity, being markedly higher in the disease-inactive state and substantially reduced during active disease. These observations indicate that dynamic regulation of polyglutamylation in IECs may be closely linked to epithelial homeostasis and inflammatory status. Although the molecular mechanisms by which polyglutamylation of IEC proteins, including the 75 kDa species, modulates epithelial barrier integrity and inflammatory responses remain to be elucidated, recent studies have expanded the repertoire of known polyglutamylated substrates beyond tubulin. For example, Dishevelled 3 (DVL3) has been identified as a polyglutamylated protein modified by TTLL family enzymes, with important consequences for protein phase separation and Wnt signaling (Kravec et al., 2024). Identification of the 75 kDa polyglutamylated protein in IECs will be a critical next step toward elucidating how polyglutamylation contributes to the regulation of epithelial barrier function and inflammatory responses in the intestinal mucosa.

SSZ has long been prescribed for IBD patients as a prodrug that is orally administered and subsequently cleaved by gut microbiota into its active anti-inflammatory component, 5-aminosalicylic acid, which is delivered to the inflamed colon. Notably, SSZ also functions as an inhibitor of xCT, but this activity is restricted to the uncleaved parent compound (Cirillo et al., 2021; Meyer et al., 2019). At the standard oral dose used in IBD therapy (4 g/day), the reported peak blood concentration of uncleaved SSZ is approximately 17.6 μg/mL (approximately 44.2 μM) (Schröder et al., 1972), a level that is unlikely to achieve robust pharmacological inhibition of xCT. In contrast, substantially higher doses of SSZ (8–12 g/day) have been administered in oncology settings to target xCT in cancer cells (Shitara et al., 2017). These observations suggest that, at clinically used doses for IBD, SSZ is unlikely to exert meaningful therapeutic effects through xCT inhibition. In our experimental setting, xCT inhibitor IKE was administered via intraperitoneal injection, thereby avoiding microbial processing in the gut and preserving their xCT-antagonistic activity. Under these conditions, IKE effectively ameliorated DSS-induced colitis, supporting a role for xCT inhibition in suppressing intestinal inflammation. Given that oral administration is preferable for patient compliance in IBD, these findings underscore the need to develop xCT inhibitors that remain stable and active within the intestinal lumen. Such agents, exemplified by IKE or related compounds, may provide a foundation for epithelial-targeted therapeutic strategies for IBD.

With respect to target suitability, xCT represents a particularly attractive candidate. Mice lacking *Slc7a11*, which encodes a functional subunit of xCT, are viable, healthy, and fertile, and do not display overt developmental or physiological abnormalities under standard conditions, suggesting that xCT is largely dispensable for organismal homeostasis (Sato et al., 2005). These observations imply that systemic inhibition of xCT may be tolerated with a relatively low risk of severe adverse effects. In addition, an increasing body of evidence indicates that xCT inhibition may confer beneficial effects in multiple contexts. For example, genetic ablation of *Slc7a11* has been reported to extend lifespan in mice, and to enhance wound healing in diabetic models compared with wild-type controls (Verbruggen et al., 2022; Maschalidi et al., 2022). Together, these findings suggest that pharmacological targeting of xCT may not only be well tolerated but could also provide ancillary benefits beyond disease-specific indications.

GS inhibition results in the retention of glutamate in tissues but also inhibits ammonia detoxification. In humans and mice, ammonia, often derived from amino acid catabolism, is detoxified by the GS and urea cycle. Mice with postnatal inducible GS deficiency in liver do not show significant increases in blood ammonia (Villar et al., 2022) whereas prenatal GS loss in the liver does show increases in blood ammonia levels (Hakvoort et al., 2017). Considering administrating GS inhibitors to human, a possible concern is that ammonia detoxification capacity may be reduced. To overcome this problem, the most critical and practical issue is to create an appropriate local drug delivery system. For example, to specifically target the IECs, pH-dependent enteric-coated tablets and extended-release formulations may be used. Our discovery that glutamate retention in IECs confers anti-inflammatory effects in colitis will provide a promising clue to development of side-effect-free therapeutics.

## Materials and Methods

### Human data set analysis

Public RNA-seq datasets (GSE186507, GSE193677, GSE206285, GSE207465) were downloaded from GEO, and corresponding raw FASTQ files were retrieved from SRA (BioProject PRJNA797175) (Argmann et al., 2023). Quantification was performed with Salmon v1.10.1 using the GRCh38 transcriptome, and gene-level TPM matrices were obtained with the tximport package in R. A total of 460 healthy control samples, 872 ulcerative colitis (UC) samples, and 1,157 Crohn’s disease (CD) samples were included in the downstream analyses.

### Mouse data set analysis

Public RNA-seq datasets (GSE131032) were downloaded from GEO, and corresponding raw FASTQ files were retrieved from SRA (BioProject PRJNA542350) (Czarnewski et al., 2019). Quantification was performed with Salmon v1.10.1 using the GRCm38 reference transcriptome, and gene-level TPM matrices were obtained with the tximport.

### Mice lines

All the mice used in this study were male on C57BL/6 background. *Slc7a11* knockout (xCT KO) mice were described (Sato al., 2005). *Slc7a11* floxed (xCT^F/F^) mice were obtained from European Mouse Mutant Archive München. Villin-Cre mice and LysM-Cre were purchased from Jackson Laboratory. All mice were housed in specific pathogen-free conditions according to the regulations of The Standards for Human Care and Use of Laboratory Animals of Tohoku University and the Guidelines for Proper Conduct of Animal Experiments by the Ministry of Education, Culture, Sports, Science, and Technology of Japan. The permission codes for the animal experiment were 2021AcA-002.

### DSS colitis model

To induce colitis, mice were given 1-3 % (w/v) dextran sulfate sodium (DSS) (MP Biomedicals, molecular weight 36-50 kDa) in the drinking water for 3-9 days (Wirtz et al., 2017). Mice were weighed every day to determine percent body weight changes, and their feces were checked daily. Disease activity index (DAI) score was calculated by the parameters, rate of weight loss, stool consistency, and degree of bloody stool (Table S1) (Wirtz et al., 2017). Colon length was measured from the proximal end of the caecum to the distal end of the rectal. A portion of dissected colon specimens was frozen for gene expression analysis, and the remaining colon was fixed in 4% paraformaldehyde. Fixed colon was embedded in paraffin and stained with haematoxylin and eosin (HE). Histological scoring was performed as previously described to assess inflammation, crypt damage, and the extent of tissue involvement (Table S2)(Dieleman, et al. 1998; Khan, et al. 2025).

L-Methionine sulfoximine (MSO) (Sigma-Aldrich, M5379) was diluted in a solution of PBS (5 mg/mL MSO). During DSS treatment, the MSO group was daily administered with 10 mg/kg/day MSO via intraperitoneal injection from day 0 to day 6 during DSS treatment. The control group was administered with an equal volume of PBS.

Imidazole ketone erastin (IKE) (Selleck chemicals, S8877) was diluted in 5% DMSO, 30% PEG300, 5% Tween80 and ddH2O (5 mg/mL IKE). During DSS treatment, the IKE group was daily administered with 25 mg/kg/day IKE (5 mL/kg/day of IKE solution) via intraperitoneal injection from day 0 to day 7 during DSS treatment. The control group was administered with an equal volume of the vehicle.

### Mucus staining

The colon paraffin sections were deparaffinized, immersed in 3% acetic acid solution for 3 minutes, and stained with 1% alcian blue staining solution at pH 1.0 (Mutoh Chemical, 4086-2) for 20 minutes. Then, the sections were rinsed twice with 3% acetic acid solution and stained with HE.

### Intestinal permeability assay

Mice were pretreated with or without 3% DSS for 4 days. Fluorescein isothiocyanate-dextran (Sigma-Aldrich, 60842-46-8) was administrated on day 4 by gavage at a dose of 440 mg/kg. The fluorescence in the plasma was quantified using a multimode plate reader (SpectraMax M2r, Molecular Devices).

### RNA purification and quantitative RT-PCR

Frozen colon tissues were homogenized using a bead homogenizer (Precellys 24, Bertin Technologies). According to the manufacturer’s instructions, total RNA samples were prepared from colon using ISOGEN (Nippon Gene, 311-02501) or ReliaPrep™ RNA Miniprep Systems (Promega, Z6011). First-strand cDNA was synthesized using ReverTra Ace qPCR RT Master Mix with gDNA Remover (TOYOBO, FSQ-301). Real-time PCR was conducted on an Applied Biosystems 7300 Real-Time PCR System (7300 M, Thermo Fisher Scientific) using THUNDERBIRD® Probe qPCR Mix (#QPS-101, TOYOBO). β-Actin was employed for normalization. All primers used for RT-PCR are described in Table S3.

### Bone marrow transplantation experiment

WT and xCT KO mice in CD45.1 background at 8-10 weeks of age were used for donors of bone marrow cells, and WT mice in CD45.2 background at 7-8 weeks of age were used for recipients. The recipient mice were given a lethal dose of radiation (7 Gy) and transplanted with 2.0 x 10^6^ bone marrow cells that were obtained from femoral bones of the donor mice via tail veins. To estimate transplant efficiency, blood samples were simultaneously collected and markers CD45.1 and CD45.2 were analyzed by flow cytometry. For surface antigen staining, Pacific Blue™-labeled anti-mouse CD45 antibody (BioLegend, 103126), FITC-labeled anti-mouse CD45.1 antibody (BD Biosciences, 553775), and PE-labeled anti-mouse CD45.2 antibody (eBioscience™, 12-0454083) were used. Propidum iodide solution (#421301, BioLegend) was added to exclude dead cells. After the staining, cells were washed and analyzed by flow cytometry (CytoFLEX LX, BECKMAN COULTER). CytExpert 2.4 (BECKMAN COULTER) was used for data analysis. After 13-15 weeks of transplantation, 3% DSS was administrated in drinking water for 9 days to induce colitis.

### Sampling feces and fecal DNA extraction for gut microbiome analysis

For gut microbiome analysis, pregnant WT mice and xCT KO mice were cohoused, and pups were in co-bred for 4 weeks from birth to weaning. After weaning, WT mice and xCT KO mice were separated in different cages for additional 16 to 18 weeks to establish the specific microbiome of each genotype before their feces were sampled. Mice were then given 1% DSS until day 6.

Isolation of fecal DNA was performed as described previously (Nagai et al., 2023). For each freeze-dried fecal sample, four 3.0 mm zirconia beads, and 100 mg of 0.1 mm zirconia/silica beads, 400 μL of DNA extraction buffer (TE containing 1% (w/v) sodium dodecyl sulfate) and 400 μL phenol/chloroform/isoamyl alcohol (25:24:1) were added and vigorously shaked (1500 rpm for 15 minutes) using a Shake Master (Biomedical Science). The resulting emulsion was centrifuged at 17,800 x □g for 10 minutes at room temperature, and the bacterial genomic DNA was purified from the aqueous phase using a standard phenol/chloroform/isoamyl alcohol protocol. RNA was digested in the sample by RNase A treatment; the resulting DNA sample then was purified again, by another round of phenol/chloroform/isoamyl alcohol treatment.

### 16S rRNA gene sequencing

16S rRNA gene sequencing was performed as described (Nagai et al., 2023). The DNA samples were analyzed using a Miseq sequencer (Illumina) to identify 16S rRNA genes in fecal microbiota. The V1–V2 region of the 16□S rRNA genes was amplified from the DNA (∼10□ng per reaction) using a universal bacterial primer set consisting of primers 27Fmod with an overhang adapter (5’-AGR GTT TGA TYM TGG CTC AG-3’) and 338R with an overhang adapter (5’-TGC TGC CTC CCG TAG GAG T-3’). PCR was performed with Tks Gflex DNA Polymerase (Takara Bio Inc), and amplification was conducted via the following program: one cycle denaturation at 98□°C for 1□min; 20 cycles of amplification at 98□°C for 10□s, 55□°C for 15□s, and 68□°C for 30□s, with final extension at 68□°C for 3□min. The amplified products were purified using Agencourt AMPure XP kits (Beckman Coulter). The purified products were then further amplified using a primer pair as follows: a forward primer (5’-AAT GAT ACG GCG ACC ACC GAG ATC TAC AC-NNNNNNNN-TAT GGT AAT TGT AGR GTT TGA TYM TGG CTC AG-3’) containing the P5 sequence, a unique 8-bp barcode sequence for each sample (indicated by the string of Ns), and an overhang adapter, as well as a reverse primer (5’-CAA GCA GAA GAC GGC ATA CGA GAT-NNNNNNNN-AGT CAG TCA GCC TGC TGC CTC CCG TAG GAG T-3’) containing the P7 sequence, a unique 8-bp barcode sequence for each sample (indicated by the string of Ns), and an overhang adapter. After purification using Agencourt AMPure XP kits, the purified products were mixed in approximately equal molar concentrations to generate a 4□nM library pool, after which the final library pool was diluted to 6 pM, including a 10% Phix Control v3 (Illumina, San Diego, California, U.S.A.) spike-in for sequencing. Finally, Miseq sequencing was performed according to the manufacturer’s instructions. In this study, 2 × 300-bp paired-end sequencing was employed. The filtered reads were processed using Quantitative Insights into Microbial Ecology 2 (QIIME2, 2019.10.0), and sequence denoising and trimming was performed using DADA2. The sequences were clustered into operational taxonomic units (OTUs) with 97% sequence homology, and OTUs were assigned to bacterial species based on classification using the RDP classifier.

### Quantification of fecal organic acids via GC/MS

The colonic organic acid concentrations were determined by gas chromatography-mass spectrometry (GC/MS) as described (Nagai et al., 2023). The frozen feces were briefly disrupted using 3 mm zirconia/silica beads (BioSpec Products) and homogenized with an extraction solution containing 100 μL of internal standard (100 μM crotonic acid), 50 μL of HCl and 200 μL ether. After vigorous shaking using Shakemaster neo (Bio Medical Science) at 1500 r.p.m. for 10 min, homogenates were centrifuged at 1000 g for 10 min, and then the top ether layer was collected and transferred to new glass vials. Aliquots (80cμL) of the ether extracts were mixed with 16□μL N-tert-butyldimethylsilyl-Nmethyltrifluoroacetamide (MTBSTFA). The vials were sealed tightly, heated at 80 °C for 20 min in a water bath, and then left at room temperature for 48 h for derivatization. The derivatized samples were run through a 6890□N Network GC System (Agilent Technologies) equipped with HP-5MS column (0.25□mm□ x □30□m x □0.25□μm) and 5973 Network Mass Selective Detector (Agilent Technologies). Pure helium (99.9999%) was used as a carrier gas and delivered at a flow rate of 1.2 mL per min. Head pressure was set at 97 kPa with a split of 20:1. The temperatures of the inlet and transfer line were 250 and 260 °C, respectively. The following temperature program was used: 60°C (3 min), 60-120°C (5°C per min), 120-300°C (20°C per min). One microliter of each sample was injected with a run time of 30□min. The concentrations of organic acids were quantified by comparing their peak areas to the standards.

### Amino acid analysis of colon tissues by liquid chromatography-electrospray ionization-tandem mass spectrometry (LC-ESI-MS/MS)

For metabolite analysis of colon tissue, WT and xCT KO mice were treated with 3% DSS for 5 days. The mice were sacrificed before or after DSS treatment and 5 mm colon tissue was taken and stored at –80℃. The frozen colon tissue was crushed with ice-cold methanol, using a bead homogenizer (Precellys 24). The sample supernatant was collected by centrifugation at 16,000 g for 5 min at 4°C. The pellet was resuspended in 0.1% SDS PBS and used to quantify total protein content by PierceTM BCA Protein Assay Kits (Thermo Fisher Scientific, 23228).

The sample supernatants were diluted to 1/5 and an isotope-labeled amino acid internal standard mixture (Sigma-Aldrich, Stable Isotope Labeled Amino Acid Mix Solution 1) was spiked into the sample and analyzed using a triple quadrupole mass spectrometry (LCMS-8060, Shimadzu Corporation) and a Nexera UHPLC system (Shimadzu Corporation). To determine the metabolites, samples were separated using a Nexera UHPLC system equipped with an Intrada AminoAcid column (Imtakt, 100 mm x 3.0 mm, 3 μm,) with the following linear gradient. Mobile phase A (0.1% formic acid in 100% acetonitrile) was dropped against mobile phase B (100 mM ammonium formate) from 100% to 75% for 4 min, 75% to 65% for 6 min, 65% to 0% for 4.33 min, and 0% was maintained for 6.67 min. The flow rate was 0.4 mL/min. The multiple reaction monitoring parameters used are listed in Table S4.

For glutamate quantification, the peak area was first normalized to that of ^13^C-labeled glutamate in the internal standard mixture and susequently normalized to the protein content of each sample. For glutamine and glutathione, ^13^C-labeled alanine and ^13^C-labeled serine were used as reference compounds, respectively, because their retention times are close to those of glutamine and glutathione. The peak areas of each analyte were normalized to the correspondoing reference compound, and calibration curves were generated based on the analyte-to-reference peak area ratios. Metabolite concentrations were calculated from these calibration curves and further normalized to the protein content of each sample.

### Preparation of mouse colon epithelial tissues

The colon was harvested from mice, thoroughly washed, and finely minced. To dissociate the epithelial layer, the colon tissue fragments were incubated in PBS containing 250 mM EDTA for 5 min. Following the incubation, the supernatant was discarded. The tissue was rinsed with PBS, and the supernatant containing detached epithelial cells was collected. The epithelial cells were pelleted by centrifugation and harvested for further analysis.

### Preparation of human UC patient biopsy specimens

Human colon biopsy samples were analyzed in this study (Figure 3C). This study was approved by the Ethics Committees of Tohoku University School of Medicine. The study was performed in accordance with the ethical standards laid down in the 1964 Declaration of Helsinki and its later amendments. This human samples and clinical information were collected through the IBD-MOCHA Study (Approval Nos. 2021-1-1244, 2023-1-106).

### Cell culture

Human colorectal cancer cell lines, Caco-2 (European Collection of Authenticated Cell Cultures) and HCT116 (American Type Culture Collection), were used. Cell lines were maintained in Dulbecco’s Modified Eagle’s Medium (DMEM) (high-glucose) with L-Glutamine and Phenol Red (Fujifilm Wako Pure Chemicals, 4429765) with 10% v/v FBS, 1% v/v penicillin/streptomycin (P/S) as glutamine containing medium (containing 4 mM glutamine) . Cells were cultured in a 5% CO_2_ incubator at 37 °C.

### Immunoblot analysis

Mouse colon epithelial tissues and human biopsy specimens were lysed in 2 x Laemmli buffer followed by boiling at 95°C for 10 min. Caco-2 cells were seeded in 6 cm culture dishes at a density of 8 × 10^5^ cells/dish. After 24 hours preincubation with normal medium (containing 4 mM glutamine), they were incubated for 24 hours with DMEM (High Glucose) with and without glutamine (Fujifilm Wako Pure Chemicals, #4429765 and #04530285) with 10% v/v FBS, 1% v/v P/S and harvested. For the preparation of whole-cell lysates, cells were directly lysed in 2 x Laemmli buffer followed by boiling at 95°C for 10 min.

The protein samples were separated by SDS-PAGE and transferred onto PVDF membranes (Immobilon P, Millipore). After the transfer of protein, the membrane was blocked in 5% (w/v) skim milk in Tris-buffered saline (TBS) containing 0.1% (w/v) Tween 20 (TBST) for 1 hour. The membranes were incubated with primary antibodies diluted with 5% (w/v) skim milk in TBST overnight at 4 °C. The antibodies used were as follows: anti polyglutamylation modification (Adipogen, AG-20B-0020B-C100) and anti-tubulin (Sigma, T9026). The membranes were washed and incubated with TBST containing peroxidase-conjugated secondary antibody for 1 hour at room temperature. The bands were detected with ECL prime (GE Healthcare) or Chemi-Lumi One L (Nacalai Tesque) using Amersham ImageQuant 800 (Cytiva). For quantitative evaluation, immunoblot band intensities were measured using Image J v1.54g software and Band/Peak Quantification Tool (Ohgane and Yoshioka, 2019).

### Transepithelial Electrical Resistance (TEER)

Caco-2 cells (2.0 × 10^5^ cells/mL) were cultured in 24-well culture incert (greiner, 662641) and used after 7 □days. Medium was changed to normal medium or without glutamine, and TEER was measured using a resistance measurement system (Thermo Fisher Scientific, Millipore). TEER was calculated as follows.

TEER (Ω x cm^2^)= (total resistance value – blank resistance value) x (effective membrane area)

### Stimulation with TNF-α

HCT116 cells were seeded in 24 well culture dishes at a density of 1 x 10^5^ cells/dish. After 24 hours preincubation with normal medium, cells were stimulated by 25 ng/mL TNF-α (Sigma-Aldrich) for 24 hours with the indicated treatment. Sulfasalazine (SSZ) was solubilized using 0.1 M NaOH and adjusting pH 8.0 using 1 M HCl (Gout et al., 2001). Total RNA was purified using ISOGEN. The gene expression was quantified by RT-PCR as described above. All primers used for RT-PCR are described in Table S3.

### Transient knockdown experiments

Transient knockdown of *TTLL4* and *TTLL5* was performed using siRNA-mediated gene silencing in the Caco-2 cell line. siGENOME SMARTpool siRNAs targeting human TTLL4 (siGENOME Human TTLL4 (9654) siRNA – SMARTpool) and TTLL5 (siGENOME Human TTLL5 (23093) siRNA – SMARTpool) were purchased from Horizon Discovery (Dharmacon). As a negative control, MISSION siRNA Universal Negative Controls (Sigma-Aldrich) were used. siCtrl and siRNAs were complexed with Lipofectamine™ RNAiMAX Transfection Reagent (Thermo Fisher Scientific) in Opti-MEM (Thermo Fisher Scientific) and transfected into cells according to the manufacturer’s instructions. The total siRNA concentration was maintained at 50 nM using siCtrl, with TTLL4 and TTLL5 each targeted at a final concentration of 25 nM. Culture media were replaced 24 h after transfection. Cells were harvested 48 h after transfection for mRNA extraction and 96 h after transfection for protein extraction to evaluate knockdown efficiency. For TEER measurements, Caco-2 cells were seeded onto transwell inserts, and siRNA transfection was performed 24 h after seeding using the same transfection protocol as described above. For TNF-α response measurements, Caco-2 cells were seeded in 6 well culture dishes at a density of 2 x 10^5^ cells/well. After 24 hours preincubation in normal medium, the siRNAs were introduced. After 3 days of transfection, cells were stimulated by 25 ng/mL TNF-α (Sigma-Aldrich) for 4 hours. Total RNA was purified using ISOGEN. The gene expression was quantified by RT-PCR as described above. All primers used for RT-PCR are described in Table S3.

### Statistical analysis

Statistical analysis between two groups was performed by Student’s *t* test or Mann-Whitney test. One way analysis of variance (one way ANOVA) was performed for three or more groups. For comparison of weight loss rates, two way ANOVA was used. Bonferroni method was used for multiple comparisons. p value < 0.05 was considered statistically significant. All the statistical analyses were performed using GraphPad Prism 10.

## Acknowledgments

We thank Drs. Naoto Ishii and Takeshi Kawabe for technical instruction of intestinal tissue analysis. We also thank the Biomedical Research Core of the IDAC Tohoku University and the Biomedical Research Core of Tohoku University Graduate School of Medicine for their technical supports.

## Funding

This work was supported by JSPS [grant numbers 25K19317 (HI), 23H02672 (HK), 22H03541(SF), 21H04799 (HM), 21H05258 (HM), 25K22535 (HM), 24H00605 (HM),], JST ERATO[grant number JPMJER1902(SF)], JST CREST [grant number JPMJCR2123(TS)], AMED [grant numbers JP21zf0127001 (HM, SF, TS), JP23gm1010009 (S.F)], the Food Science Institute Foundation (SF) and Human Biology-Microbiome-Quantum Research Center (WPI-Bio2Q), Keio University (TS). The funders had no role in the study design, data collection and analysis, decision to publish or manuscript preparation.

## Author contributions

H.I., H.Sekine. and H.M. designed the research and conducted the experiments. Y.Y., N.K. and H.K. performed cell and animal experiments. H.Sato. provided a critical material for experiments. H.H., S.M., F.-Y.W., S.F. and T.S. conducted microbiome and metabolome analysis. T.K., Y.K., and A.M. analyzed the data. A.M., H.Sekine. and H.M. supervised the research. H.I., H.Sekine. and H.M. wrote an early draft of the paper.

## Competing interests

The authors declare no competing financial or nonfinancial interests.

## Data and materials availability

All data are available in the main text or the supplementary materials.

## Notes

### Competing Interest Statement

The authors have declared no competing interest.

## References

Alatab S., et al., GBD 2017 Inflammatory Bowel Disease Collaborators. The global, regional, and national burden of inflammatory bowel disease in 195 countries and territories, 1990-2017: a systematic analysis for the Global Burden of Disease Study 2017. Lancet Gastroenterol Hepatol. 2020 Jan;5(1):17–30. doi: 10.1016/S2468-1253(19)30333-4.

Argmann C, Hou R, Ungaro RC, Irizar H, Al-Taie Z, Huang R, Kosoy R, Venkat S, Song WM, Di’Narzo AF, Losic B, Hao K, Peters L, Comella PH, Wei G, Atreja A, Mahajan M, Iuga A, Desai PT, Branigan P, Stojmirovic A, Perrigoue J, Brodmerkel C, Curran M, Friedman JR, Hart A, Lamousé-Smith E, Wehkamp J, Mehandru S, Schadt EE, Sands BE, Dubinsky MC, Colombel JF, Kasarskis A, Suárez-Fariñas M. Biopsy and blood-based molecular biomarker of inflammation in IBD. Gut. 2023 Jul;72(7):1271–1287. doi: 10.1136/gutjnl-2021-326451.

Brinkman BM, Becker A, Ayiseh RB, Hildebrand F, Raes J, Huys G, Vandenabeele P. Gut microbiota affects sensitivity to acute DSS-induced colitis independently of host genotype. Inflamm Bowel Dis. 2013 Nov;19(12):2560–7. doi: 10.1097/MIB.0b013e3182a8759a.

Chang JT. Pathophysiology of Inflammatory Bowel Diseases. N Engl J Med. 2020 Dec 31;383(27):2652–2664. doi: 10.1056/NEJMra2002697.

Chassaing B, Aitken JD, Malleshappa M, Vijay-Kumar M. Dextran sulfate sodium (DSS)-induced colitis in mice. Curr Protoc Immunol. 2014 Feb 4;104:15.25.1–15.25.14. doi: 10.1002/0471142735.im1525s104.

Cirillo D, Sarowar S, Øyvind Enger P, Bjørsvik HR. Structure-Activity-Relationship-Aided Design and Synthesis of xCT Antiporter Inhibitors. ChemMedChem. 2021 Sep 6;16(17):2650–2668. doi: 10.1002/cmdc.202100204. Epub 2021 May 28. PMID: 33847044; PMCID: PMC8518981.

Cosnes J, Gower-Rousseau C, Seksik P, Cortot A. Epidemiology and natural history of inflammatory bowel diseases. Gastroenterology. 2011 May;140(6):1785–94. doi: 10.1053/j.gastro.2011.01.055.

Czarnewski P, Parigi SM, Sorini C, Diaz OE, Das S, Gagliani N, Villablanca EJ. Conserved transcriptomic profile between mouse and human colitis allows unsupervised patient stratification. Nat Commun. 2019 Jun 28;10(1):2892. doi: 10.1038/s41467-019-10769-x.

Dawson PA, Huxley S, Gardiner B, Tran T, McAuley JL, Grimmond S, McGuckin MA, Markovich D. Reduced mucin sulfonation and impaired intestinal barrier function in the hyposulfataemic NaS1 null mouse. Gut. 2009 Jul;58(7):910–9. doi: 10.1136/gut.2007.147595.

Dieleman LA, Palmen MJ, Akol H, Bloemena E, Peña AS, Meuwissen SG, Van Rees EP. Chronic experimental colitis induced by dextran sulphate sodium (DSS) is characterized by Th1 and Th2 cytokines. Clin Exp Immunol. 1998 Dec;114(3):385–91. doi: 10.1046/j.1365-2249.1998.00728.x.

Eddé B, Rossier J, Le Caer JP, Desbruyères E, Gros F, Denoulet P. Posttranslational glutamylation of alpha-tubulin. Science. 1990 Jan 5;247(4938):83–5. doi: 10.1126/science.1967194.

Eichele DD, Kharbanda KK. Dextran sodium sulfate colitis murine model: An indispensable tool for advancing our understanding of inflammatory bowel diseases pathogenesis. World J Gastroenterol. 2017 Sep 7;23(33):6016–6029. doi: 10.3748/wjg.v23.i33.6016.

Forster SC, Clare S, Beresford-Jones BS, Harcourt K, Notley G, Stares MD, Kumar N, Soderholm AT, Adoum A, Wong H, Morón B, Brandt C, Dougan G, Adams DJ, Maloy KJ, Pedicord VA, Lawley TD. Identification of gut microbial species linked with disease variability in a widely used mouse model of colitis. Nat Microbiol. 2022 Apr;7(4):590–599. doi: 10.1038/s41564-022-01094-z.

Friedrich M, Diegelmann J, Beigel F, Brand S. IL-17A alone weakly affects the transcriptome of intestinal epithelial cells but strongly modulates the TNF-α-induced expression of inflammatory mediators and inflammatory bowel disease susceptibility genes. Inflamm Bowel Dis. 2014 Sep;20(9):1502–15. doi: 10.1097/MIB.0000000000000121.

Gout PW, Buckley AR, Simms CR, Bruchovsky N. Sulfasalazine, a potent suppressor of lymphoma growth by inhibition of the x(c)- cystine transporter: a new action for an old drug. Leukemia. 2001 Oct;15(10):1633–40. doi: 10.1038/sj.leu.2402238.

Häberle J, Görg B, Rutsch F, Schmidt E, Toutain A, Benoist JF, Gelot A, Suc AL, Höhne W, Schliess F, Häussinger D, Koch HG. Congenital glutamine deficiency with glutamine synthetase mutations. N Engl J Med. 2005 Nov 3;353(18):1926–33. doi: 10.1056/NEJMoa050456.

Hakvoort TB, He Y, Kulik W, Vermeulen JL, Duijst S, Ruijter JM, Runge JH, Deutz NE, Koehler SE, Lamers WH. Pivotal role of glutamine synthetase in ammonia detoxification. Hepatology. 2017 Jan;65(1):281–293. doi: 10.1002/hep.28852.

He Y, Hakvoort TB, Vermeulen JL, Labruyère WT, De Waart DR, Van Der Hel WS, Ruijter JM, Uylings HB, Lamers WH. Glutamine synthetase deficiency in murine astrocytes results in neonatal death. Glia. 2010 Apr 15;58(6):741–54. doi: 10.1002/glia.20960.

Khan MJ, Lee YJ, Lee SY, Chung H, Nguyen-Phuong T, Kim YH, Park CG, Kang YM. Novel Autologous Regulatory T-Cell Therapy Ameliorates DSS-Induced Colitis in Humanized Mice. Inflamm Bowel Dis. 2025 Sep 1;31(9):2535–2546. doi: 10.1093/ibd/izaf141.

Kiesler P, Fuss IJ, Strober W. Experimental Models of Inflammatory Bowel Diseases. Cell Mol Gastroenterol Hepatol. 2015 Mar 1;1(2):154–170. doi: 10.1016/j.jcmgh.2015.01.006.

Kravec M, Šedo O, Nedvědová J, Micka M, Šulcová M, Zezula N, Gömöryová K, Potěšil D, Sri Ganji R, Bologna S, Červenka I, Zdráhal Z, Harnoš J, Tripsianes K, Janke C, Bařinka C, Bryja V. Carboxy-terminal polyglutamylation regulates signaling and phase separation of the Dishevelled protein. EMBO J. 2024 Nov;43(22):5635–5666. doi: 10.1038/s44318-024-00254-7.

Lamb CA, Kennedy NA, Raine T, Hendy PA, Smith PJ, Limdi JK, Hayee B, Lomer MCE, Parkes GC, Selinger C, Barrett KJ, Davies RJ, Bennett C, Gittens S, Dunlop MG, Faiz O, Fraser A, Garrick V, Johnston PD, Parkes M, Sanderson J, Terry H; IBD guidelines eDelphi consensus group; Gaya DR, Iqbal TH, Taylor SA, Smith M, Brookes M, Hansen R, Hawthorne AB. British Society of Gastroenterology consensus guidelines on the management of inflammatory bowel disease in adults. Gut. 2019 Dec;68(Suppl 3):s1–s106. doi: 10.1136/gutjnl-2019-318484. Epub 2019 Sep 27. Erratum in: Gut. 2021 Apr;70(4):1. doi: 10.1136/gutjnl-2019-318484corr1.

Laukens D, Brinkman BM, Raes J, De Vos M, Vandenabeele P. Heterogeneity of the gut microbiome in mice: guidelines for optimizing experimental design. FEMS Microbiol Rev. 2016 Jan;40(1):117–32. doi: 10.1093/femsre/fuv036.

Louis P, Flint HJ. Formation of propionate and butyrate by the human colonic microbiota. Environ Microbiol. 2017 Jan;19(1):29–41. doi: 10.1111/1462-2920.13589.

Madison BB, Dunbar L, Qiao XT, Braunstein K, Braunstein E, Gumucio DL. Cis elements of the villin gene control expression in restricted domains of the vertical (crypt) and horizontal (duodenum, cecum) axes of the intestine. J Biol Chem. 2002 Sep 6;277(36):33275–83. doi: 10.1074/jbc.M204935200.

Maschalidi S, Mehrotra P, Keçeli BN, De Cleene HKL, Lecomte K, Van der Cruyssen R, Janssen P, Pinney J, van Loo G, Elewaut D, Massie A, Hoste E, Ravichandran KS. Targeting SLC7A11 improves efferocytosis by dendritic cells and wound healing in diabetes. Nature. 2022 Jun;606(7915):776–784. doi: 10.1038/s41586-022-04754-6.

Matsuoka K, Kobayashi T, Ueno F, Matsui T, Hirai F, Inoue N, Kato J, Kobayashi K, Kobayashi K, Koganei K, Kunisaki R, Motoya S, Nagahori M, Nakase H, Omata F, Saruta M, Watanabe T, Tanaka T, Kanai T, Noguchi Y, Takahashi KI, Watanabe K, Hibi T, Suzuki Y, Watanabe M, Sugano K, Shimosegawa T. Evidence-based clinical practice guidelines for inflammatory bowel disease. J Gastroenterol. 2018 Mar;53(3):305–353. doi: 10.1007/s00535-018-1439-1.

Meyer AR, Engevik AC, Willet SG, Williams JA, Zou Y, Massion PP, Mills JC, Choi E, Goldenring JR. Cystine/Glutamate Antiporter (xCT) Is Required for Chief Cell Plasticity After Gastric Injury. Cell Mol Gastroenterol Hepatol. 2019;8(3):379–405. doi: 10.1016/j.jcmgh.2019.04.015. Epub 2019 May 6. PMID: 31071489; PMCID: PMC6713894.

Nagai M, Moriyama M, Ishii C, Mori H, Watanabe H, Nakahara T, Yamada T, Ishikawa D, Ishikawa T, Hirayama A, Kimura I, Nagahara A, Naito T, Fukuda S, Ichinohe T. High body temperature increases gut microbiota-dependent host resistance to influenza A virus and SARS-CoV-2 infection. Nat Commun. 2023 Jun 30;14(1):3863. doi: 10.1038/s41467-023-39569-0.

Ohgane K & Yoshioka H. Quantification of gel bands by an Image J macro, band/peak quantification tool. protocols.io 2019. doi.org/ 10.17504/protocols.io.7vghn3w.

Pan X, Zhu Q, Pan LL, Sun J. Macrophage immunometabolism in inflammatory bowel diseases: From pathogenesis to therapy. Pharmacol Ther. 2022 Oct;238:108176. doi: 10.1016/j.pharmthera.2022.108176.

Parada Venegas D, De la Fuente MK, Landskron G, González MJ, Quera R, Dijkstra G, Harmsen HJM, Faber KN, Hermoso MA. Short Chain Fatty Acids (SCFAs)-Mediated Gut Epithelial and Immune Regulation and Its Relevance for Inflammatory Bowel Diseases. Front Immunol. 2019 Mar 11;10:277. doi: 10.3389/fimmu.2019.00277.

Parker JL, Deme JC, Kolokouris D, Kuteyi G, Biggin PC, Lea SM, Newstead S. Molecular basis for redox control by the human cystine/glutamate antiporter system xc. Nat Commun. 2021 Dec 8;12(1):7147. doi: 10.1038/s41467-021-27414-1.

Rüdiger M, Plessman U, Klöppel KD, Wehland J, Weber K. Class II tubulin, the major brain beta tubulin isotype is polyglutamylated on glutamic acid residue 435. FEBS Lett. 1992 Aug 10;308(1):101–5. doi: 10.1016/0014-5793(92)81061-p.

Ruse CI, Chin HG, Pradhan S. Polyglutamylation: biology and analysis. Amino Acids. 2022 Apr;54(4):529–542. doi: 10.1007/s00726-022-03146-4.

Sato H, Shiiya A, Kimata M, Maebara K, Tamba M, Sakakura Y, Makino N, Sugiyama F, Yagami K, Moriguchi T, Takahashi S, Bannai S. Redox imbalance in cystine/glutamate transporter-deficient mice. J Biol Chem. 2005 Nov 11;280(45):37423–9. doi: 10.1074/jbc.M506439200.

Schröder H, Evans DA. Acetylator phenotype and adverse effects of sulphasalazine in healthy subjects. Gut. 1972 Apr;13(4):278–84. doi: 10.1136/gut.13.4.278.

Shiomi Y, Nishiumi S, Ooi M, Hatano N, Shinohara M, Yoshie T, Kondo Y, Furumatsu K, Shiomi H, Kutsumi H, Azuma T, Yoshida M. GCMS-based metabolomic study in mice with colitis induced by dextran sulfate sodium. Inflamm Bowel Dis. 2011 Nov;17(11):2261–74. doi: 10.1002/ibd.21616.

Shitara K, Doi T, Nagano O, Imamura CK, Ozeki T, Ishii Y, Tsuchihashi K, Takahashi S, Nakajima TE, Hironaka S, Fukutani M, Hasegawa H, Nomura S, Sato A, Einaga Y, Kuwata T, Saya H, Ohtsu A. Dose-escalation study for the targeting of CD44v^+^ cancer stem cells by sulfasalazine in patients with advanced gastric cancer (EPOC1205). Gastric Cancer. 2017 Mar;20(2):341–349. doi: 10.1007/s10120-016-0610-8.

Torrino S, Grasset EM, Audebert S, Belhadj I, Lacoux C, Haynes M, Pisano S, Abélanet S, Brau F, Chan SY, Mari B, Oldham WM, Ewald AJ, Bertero T. Mechano-induced cell metabolism promotes microtubule glutamylation to force metastasis. Cell Metab. 2021 Jul 6;33(7):1342–1357.e10. doi: 10.1016/j.cmet.2021.05.009.

Van De Walle J, Hendrickx A, Romier B, Larondelle Y, Schneider YJ. Inflammatory parameters in Caco-2 cells: effect of stimuli nature, concentration, combination and cell differentiation. Toxicol In Vitro. 2010 Aug;24(5):1441–9. doi: 10.1016/j.tiv.2010.04.002.

Verbruggen L, Ates G, Lara O, De Munck J, Villers A, De Pauw L, Ottestad-Hansen S, Kobayashi S, Beckers P, Janssen P, Sato H, Zhou Y, Hermans E, Njemini R, Arckens L, Danbolt NC, De Bundel D, Aerts JL, Barbé K, Guillaume B, Ris L, Bentea E, Massie A. Lifespan extension with preservation of hippocampal function in aged system x_c_^-^-deficient male mice. Mol Psychiatry. 2022 Apr;27(4):2355–2368. doi: 10.1038/s41380-022-01470-5.

Villar VH, Allega MF, Deshmukh R, Ackermann T, Nakasone MA, Vande Voorde J, Drake TM, Oetjen J, Bloom A, Nixon C, Müller M, May S, Tan EH, Vereecke L, Jans M, Blancke G, Murphy DJ, Huang DT, Lewis DY, Bird TG, Sansom OJ, Blyth K, Sumpton D, Tardito S. Hepatic glutamine synthetase controls N^5^-methylglutamine in homeostasis and cancer. Nat Chem Biol. 2023 Mar;19(3):292–300. doi: 10.1038/s41589-022-01154-9.

Wirtz S, Popp V, Kindermann M, Gerlach K, Weigmann B, Fichtner-Feigl S, Neurath MF. Chemically induced mouse models of acute and chronic intestinal inflammation. Nat Protoc. 2017 Jul;12(7):1295–1309. doi: 10.1038/nprot.2017.044.

