## Supplementary material for "Epithelial Glutamate Retention Protects against Colitis": Figure S1, S2, and Table S1-S4

†**Corresponding authors:**

Hiroki Sekine, Ph.D.

**The PDF file includes:**

Figs. S1 to S2

Tables S1 to S4

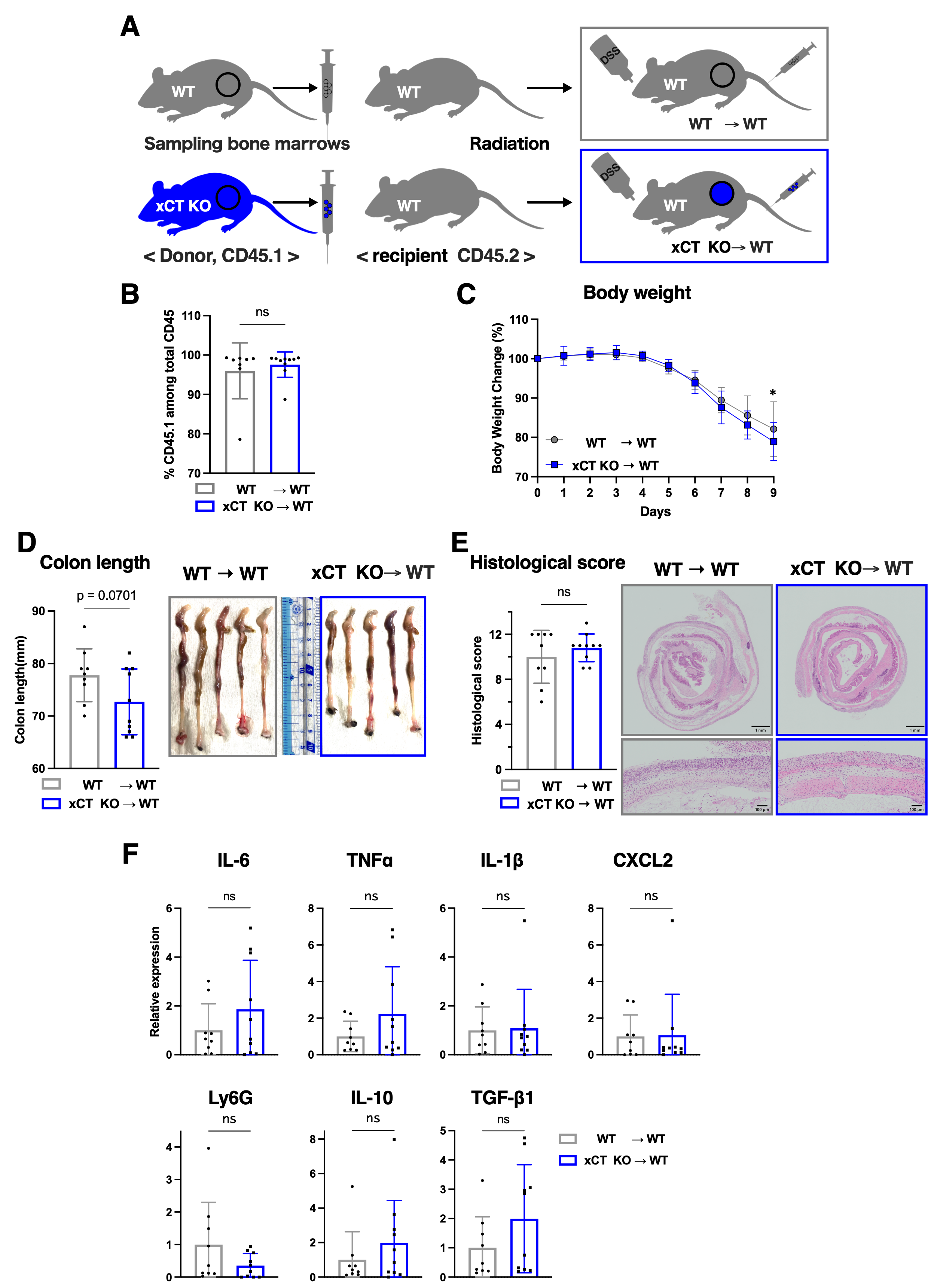

**Fig. S1. Transplantation of xCT knockout bone marrow cells failed to ameliorate DSS-induced colitis in recipient mice.**

**A.** Scheme of bone marrow transplantation experiments and DSS colitis. The bone marrow cells were collected from WT and xCT KO donor mice (CD45.1) and transplanted to lethally irradiated WT recipient mice (CD45.2). Recipient mice transplanted with WT bone marrow were described as WT →WT, and those with xCT KO bone marrow were described as xCT KO →WT.

**B.** Chimerism analysis for evaluation of transplantation efficiency. The mean percentage of CD45.1 in blood samples was 95.99% for WT→WT mice (n=8) and 97.99% for xCT KO→WT mice (n=10). Data are shown as mean ± SD from two independent experiments.

**C-E.** WT→WT mice (n=9) and xCT KO→WT mice (n=10) were treated with 3% DSS at 13 -15 weeks after the transplantation. Body weight changes **(C)** and colon length measurement and macroscopic observation of colon tissues **(D)** are shown. Histological score and HE staining of colon tissues are shown with scale bars corresponding to 1 mm for lower magnification and 100 μm for higher magnification (**E**). Data are shown as mean ± SD from two independent experiments.

**F.** RT-PCR for measuring mRNA expression of cytokines, chemokines and a neutrophil marker, Ly6G, in their colon tissues with 3% DSS treatment on day 9 (WT mice; n=9, xCT KO→WT mice; n=10). β-Actin was employed for normalization. Mean values of WT→WT mice were set as 1. Data are shown as mean ± SD from two independent experiments.

Data represent means ± standard deviation. **B** and **D**-**F** were analyzed by two-sided Student's t-test. **C** was analyzed by two-way ANOVA. ns, not significant.

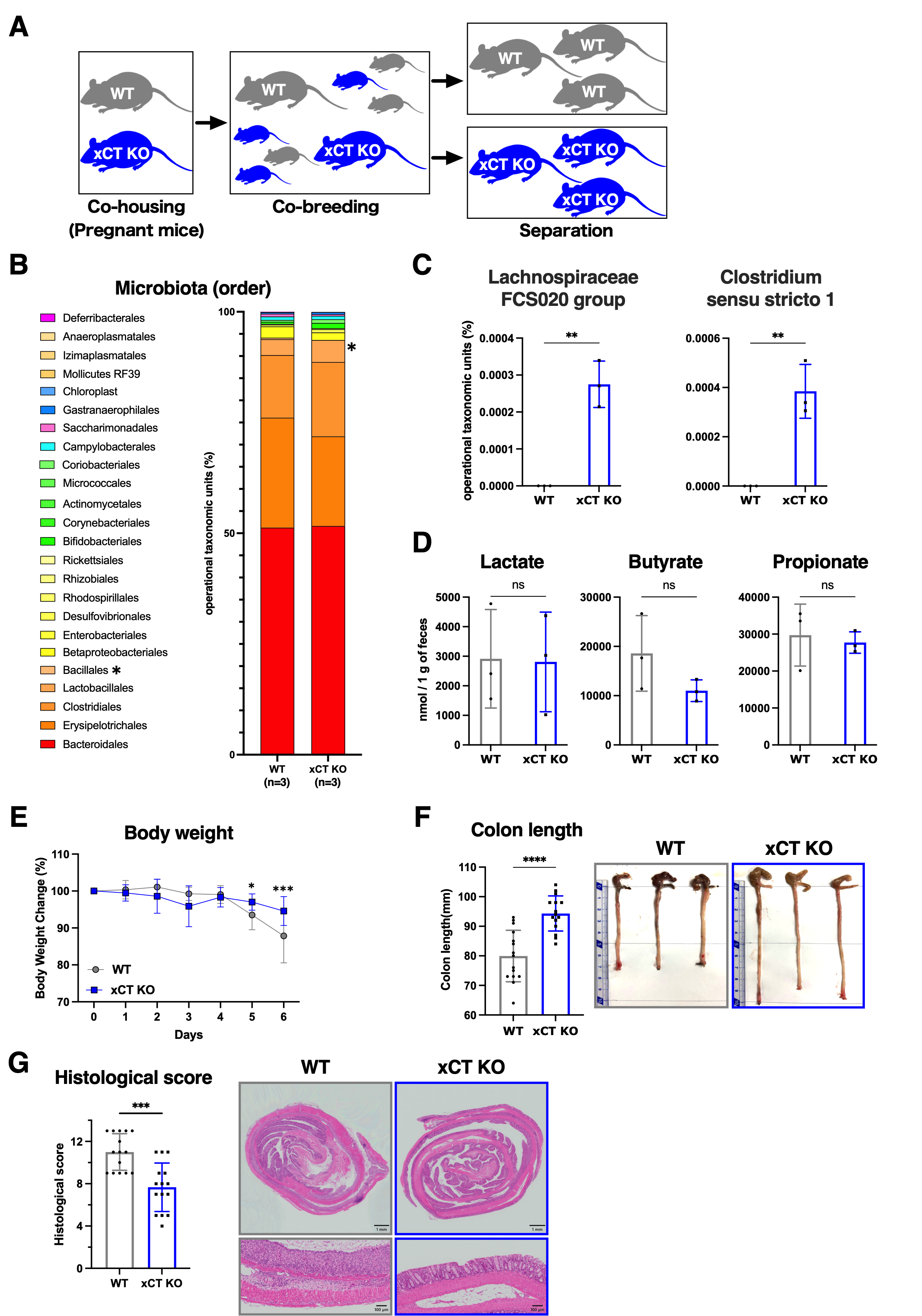

**Fig. S2. Co-breeding does not cancel anti-inflammatory effects of xCT deficiency.**

**A.** Scheme of gut microbiota and metabolite analysis. To equalize the gut microbiota, WT and xCT KO pregnant mice were co-housed, and the pups were co-bred for 4 weeks from birth to weaning. Then, the mice were separated by genotype. Additional 16-18 weeks later, their feces were collected, and DSS treatment was initiated.

**B**-**D.** Gut microbiota analysis. (WT mice; n=3, xCT KO mice; n=3) Gut microbiota diversity at the order level **(B)**, gut commensal bacteria at the genus level increased in xCT KO mice **(C)** and quantification of SCFAs in fecal metabolites **(D)**.

**E**-**G.** The mice whose feces were analyzed were treated with 1% DSS to induce colitis. Body weight changes **(E)** and colon length measurement and macroscopic observation of colon tissues **(F)**. Histological score and HE staining of colon tissues are shown with scale bars corresponding to 1 mm for lower magnification and 100 μm for higher magnification (**G**). Data are shown as mean ± SD from three independent experiments.

Data represent means ± standard deviation. **B**-**D**, **F** and **G** were analyzed by two-sided Student's t-test. **E** was analyzed by two-way ANOVA. *p<0.05, **p<0.01, ***p<0.001,****p<0.0001; ns, not significant.

**Table S1. Disease activity index (DAI) score of DSS colitis model.**

| Score | Body weight decrease | Stool consistency | Rectal bleeding |
| --- | --- | --- | --- |
| 0 | 〜 < 1% | normal | negative hemoccult |
| 1 | 1% ≦ 〜 ＜ 6% | soft but still formed | negative hemoccult |
| 2 | 6% ≦ 〜 ＜ 11% | soft | positive hemoccult |
| 3 | 11% ≦ 〜 ＜ 18% | very soft; wet | blood traces in stool visible |
| 4 | 18% ≦ 〜 | watery diarrhea | gross rectal bleeding |

Disease activity index (DAI) score: sum of the scores of 3 parameters (0-12)

**Table S2. Histological score of DSS colitis model.**

| Score | Inflammation | Extent or depth of injury | Crypt damage | Percentage (%) of tissue involvement |
| --- | --- | --- | --- | --- |
| 0 | None | None | No Damage | None |
| 1 | Slight (Small, focal, or widely separated, limited to lamina propria) | Mucosal | Basal 1/3 Damage (Loss of bottom one-third of the crypts) | 1-25% (Up to 25% of the tissue effected by disease process) |
| 2 | Moderate (Multifocal or locally extensive, extending to submucosa) | Mucosal and submucosal (Mural) | Basal 2/3 Damage  (Loss of bottom two-third of the crypts) | 26-50%  (Up to 50% of the tissue effected by disease process) |
| 3 | Severe (Transmural inflammation with ulcers covering >20 crypts) | Transmural  (Involve all layer of gut) | Only surface epithelium intact (Loss of entire crypt with the surface epithelium remaining intact) | 51-75%  (Up to 75% of the tissue effected by disease process) |
| 4 | - | - | Entire crypt and epithelium lost  (Loss of the entire crypt and surface epithelium) | 76-100%  (More Than 75% of the tissue effected by disease process) |

| Gene |  | Sequence |
| --- | --- | --- |
| Mouse β-Actin | Forward | 5'-CGGTTCCGATGCCCTGAGGCTCTT-3' |
|  | Reverse | 5'-CGTCACACTTCATGATGGAATTGA-3' |
| Mouse IL-6 | Forward | 5'-CTGCAAGAGACTTCCATCCAG-3' |
|  | Reverse | 5'-AGTGGTATAGACAGGTCTGTTGG-3' |
| Mouse Tnfα | Forward | 5'-CACGCTCTTCTGTCTACTGAA-3' |
|  | Reverse | 5'-GGCTACAGGCTTGTCACTCGA-3' |
| Mouse IL-1β | Forward | 5'-TGCCACCTTTTGACAGTGATG-3' |
|  | Reverse | 5'-TGATGTGCTGCTGCGAGATT-3' |
| Mouse Cxcl2 | Forward | 5'-TCCAGAGCTTGAGTGTGACG-3' |
|  | Reverse | 5'-TAGCCTTGCCTTTGTTCAGT-3' |
| Mouse Ly6G | Forward | 5'-GGGCTGGAGTGCTACAATTG-3' |
|  | Reverse | 5'-GCAGATGGGAAGGCAGAGAT-3' |
| Mouse IL-10 | Forward | 5'-GGCGCTGTCATCGATTTCTC-3' |
|  | Reverse | 5'-ATGGCCTTGTAGACACCTTGG-3' |
| Mouse Tgfβ | Forward | 5'-CAGCAGCCGGTTACCAAG-3' |
|  | Reverse | 5'-TGGAGCAACATGTGGAACTC-3' |
| Mouse Slc7a11（xCT） | Forward | 5'-TGGGTGGAACTGCTCGTAAT-3' |
|  | Reverse | 5'-AGGATGTAGCGTCCAAATGC-3' |
| Human β-ACTIN | Forward | 5'-TGGCACCCAGCACAATGAA-3' |
|  | Reverse | 5'-CTAAGTCATAGTCCGCCTAGAAGCA-3' |
| Human IL-8 | Forward | 5'-CTCTTGGCAGCCTTCCTGATTTCT-3' |
|  | Reverse | 5'-GGGTGGAAAGGTTTGGAGTATGTCT-3' |
| Human TTLL4 | Forward | 5'-gacagtgaggacaccagcaa-3' |
|  | Reverse | 5'-ctggaaagtggaggaagcag-3' |
| Human TTLL5 | Forward | 5'-cacccaagcagcactgacta-3' |
|  | Reverse | 5'-cgcagtaagaagggcttcag-3' |

**Table S3. Primers for qPCR.**

**Table S4. Multiple reaction monitoring parameters of metabolites used for LC-ESI-MS/MS analyses.**

| **Analytes** | **Polarity** | **Precursor ion (m/z)** | **Product ion (m/z)** | **Collision energy (V)** |
| --- | --- | --- | --- | --- |
| Glutamate | + | 148.10 | 102.05 | –14.0 |
| Glutamate-^13^C_5_ | + | 153.10 | 106.05 | –14.0 |
| Glutamine | + | 147.00 | 84.05 | –20.0 |
| Alanine-^13^C_3_^*^ | + | 94.10 | 48.1 | –13.0 |
| Glutathione | + | 308.10 | 179.10 | –11.0 |
| Serine-^13^C_3_^*^ | + | 110.10 | 63.10 | –13.0 |

^*^Alanine-^13^C_3_ and serine-^13^C_3_ were used as reference compounds for glutamine and glutathione, respectively.
